# De novo extraction of microbial strains from metagenomes reveals intra-species niche partitioning

**DOI:** 10.1101/073825

**Authors:** Christopher Quince, Stephanie Connelly, Sébastien Raguideau, Johannes Alneberg, Seung Gu Shin, Gavin Collins, A. Murat Eren

## Abstract

**Background:** We introduce `DESMAN` for De novo Extraction of Strains from MetAgeNomes. Metagenome sequencing generates short reads from throughout the genomes of a microbial community. Increasingly large, multi-sample metagenomes, stratified in space and time are being generated from communities with thousands of species. Repeats result in fragmentary co-assemblies with potentially millions of contigs. Contigs can be binned into metagenome assembled genomes (MAGs) but strain level variation will remain. DESMAN identifies variants on core genes, then uses co-occurrence across samples to link variants into strain sequences and abundance profiles. These strain profiles are then searched for on non-core genes to determine the accessory genes present in each strain.

**Results:** We validated `DESMAN` on a synthetic twenty genome community with 64 samples. We could resolve the five *E. coli* strains present with 99.58% accuracy across core gene variable sites and their gene complement with 95.7% accuracy. Similarly, on real fecal metagenomes from the 2011 *E. coli* (STEC) O104:H4 outbreak, the outbreak strain was reconstructed with 99.8% core sequence accuracy. Application to an anaerobic digester metagenome time series reveals that strain level variation is endemic with 16 out of 26 MAGs (61.5%) examined exhibiting two strains. In almost all cases the strain proportions were not statistically different between replicate reactors, suggesting intra-species niche partitioning. The only exception being when the two strains had almost identical gene complement and, hence, functional capability.

**Conclusions:** `DESMAN` will provide a provide a powerful tool for *de novo* resolution of fine-scale variation in microbial communities. It is available as open source software from https://github.com/chrisquince/DESMAN.

## Background

Metagenomics, the direct sequencing of DNA extracted from an environment, offers an unique opportunity to study whole microbial communities *in situ*. The majority of contemporary metagenomics studies use shotgun sequencing, where DNA is fragmented prior to sequencing with short reads, of the order of hundreds of base pairs (bps). To fully realise the potential of metagenomics, methods capable of resolving both the species and the strains present from this data are needed. There exist reference-based solutions for strain identification [36, 35]. However, for the vast majority of microbial species comprehensive strain-level databases do not exist. A situation that is unlikely to change, particularly given the great diversity of unculturable microbes [7]. This motivates *de novo* strategies capable of resolving novel variation at high resolution direct from metagenome data.

It is not possible to simply assemble the reads into the strain genomes present. This is because in the presence of repeats, identical regions that exceed the read length, assemblies become uncertain and fragment into multiple contigs [32]. Metagenomes contain many repeats precisely because of strain variation and hence produce highly fragmented assemblies. It is possible to ‘bin’ these contigs into partitions that derive from the same species using sequence composition [20, 38] and more powerfully the varying coverage of individual contigs over multiple samples [37, 2, 3, 13] but the resulting resulting genome bins, or ‘metagenome-assembled genomes’ (MAGs), represent aggregates of multiple similar strains. These strains will vary both in the precise sequence of shared genes, when that variation is below the resolution of the assembler, but also in gene complement, not all genes and, hence, contigs will be present in all strains.

Recently, two methods have been developed which use the frequency of variants across multiple samples, when mapped against reference genes or genomes to *de novo* resolve strain level variation, these are the `Lineage` algorithm of O’Brien *et al.* [30], and `constrains` [27]. Crucially though, no method has yet been developed that works from assembled contigs avoiding the need for any reference genomes, and, hence, is applicable to unculturable organisms. Here, we show that it is possible to combine this principle with contig binning algorithms and resolve the strain level variation in MAGs both nucleotide variation on core genes and variation in gene complement.

We denote our strategy, `DESMAN` - De novo Extraction of Strains from MetAgeNomes. We assume that a coassembly has been performed and contigs binned into MAGs, any binning algorithm could be used for this but here we applied `CONCOCT` [3]. We also assume that reads have been mapped back onto these contigs as part of this process. To resolve strain variation within a MAG or group of MAGs deriving from a single species, we first identify core genes, genes present in all strains in a single copy. In the absence of any reference genomes these will simply be those genes known to be core for all bacteria and archaea e.g. the 36 Clusters of Orthologous Groups of proteins (COGs) identified in [3]. If reference genomes from the same species or related taxa are available then these can be used to identify further genes that will satisfy the criteria of being present in all strains in a single copy, in which case we denote these single-copy core species genes (SCSGs). Using the read mappings we calculate the base frequencies at each position on the SCSGs. Next we determine variant positions using a likelihood ratio test applied to the frequencies of each base summed across samples. We then use the base frequencies across samples on these variant positions to resolve the number of strains present, their abundance and their unique sequence or haplotype at each variant position for each SCSG.

The second component of `DESMAN` is to use this information to determine which accessory genes are present in which strain. From the analysis of SCSGs we know how many strains are present and their relative abundances across samples. The signature of relative frequencies across samples associated with each strain will also be observed on the non-core gene variants but, crucially, not all strains will possess these genes and potentially they may be in multiple-copies. The strain relative frequencies have to be adjusted therefore to reflect these copy numbers. For instance, if a gene is present in just a single copy in one strain it can have no variants. In addition, the total coverage associated with a gene will also depend on which strains possess that gene being a simple sum of the individual strain coverages. Here, we do not address the multi-copy problem, just gene presence or absence in a strain. We infer these given the observed variant base frequencies and gene coverages across samples whilst keeping the strain signatures fixed at those computed from the SCSGs. This also provides a strategy for inferring non-core gene haplotypes on strains. Taken together these two steps provide a procedure for resolving both strain haplotypes on the core genome and their gene complements entirely *de novo* from short read metagenome data. We recommend applying this strategy to genes, but crucially genes called on the assembled contigs. If contig assignments are preferred, the same methodology could be applied directly to the contigs themselves, or a consensus assignment of genes on a contig used to determine presence or absence in a given strain.

The advantage of using base frequencies across samples to resolve strains, rather than existing haplotype resolution algorithms that link variants using reads [40], is that it enables us to resolve variation that is less divergent than the reciprocal of the read length and link strains across contigs. The intuition behind frequency based strain inference is similar to that of contig binning, the frequencies of variants associated with a strain fluctuate across samples with the abundance of that strain. However, in this case it is necessary to account for the fact that multiple strains may share the same nucleotide identity at a given variant position. To solve this problem we develop a full Bayesian model, fit by a Markov Chain Monte Carlo (MCMC) Gibbs sampler, for the strain frequencies, their haplotypes and also sequencing error rates. To improve convergence we initialise the Gibbs sampler using non-negative matrix factorisation, or more properly non-negative tensor factorisation (NTF), a method from machine learning that is equivalent to the maximum likelihood solution [39]. Our approach is very similar to the `Lineage` algorithm developed by O’Brien *et al.* [30], except that they have a simpler noise model but a more complex prior for the strain haplotypes derived from an underlying phylogenetic tree. Both our approaches differ from the heuristic strategy for strain inference used in constrains[27]. The full Bayesian approach allows not just a single estimate of the strain haplotypes, but also an estimate of the uncertainty in the predictions through comparison of replicate MCMC runs.

To illustrate the efficacy of the `DESMAN` pipeline we first apply it to the problem of resolving *Escherichia coli* strains in metagenomic data sets. *E. coli* has a highly variable genome [18], and while some strains of *E. coli* occur as harmless commensals in the human gut, others can be harmful pathogens. We used a synthetic data set of 64 samples generated from an *in sillico* community comprising 5*E. coli* strains, and 15 other strains commonly found in human gut samples. Strains in this data set were present in each sample with varying abundances determined by the 16S rRNA community profiles we obtained from the HMP project [16]. The reads themselves simulated a typical HiSeq 2500 run. We then used 53 real fecal metagenomes generated from the 2011 Shiga toxin-producing *E. coli* (STEC) O104:H4 outbreak samples [25] to validate our ability to correctly resolve the outbreak strain using `DESMAN`. The results from these analyses were encouraging but the real potential of `DESMAN` is to resolve strains for environmental, potentially unculturable organisms. To demonstrate this we applied the pipeline to samples taken from a time series of three replicate anaerobic digestion reactors, successfully resolving strain variation in 16 different MAGs including some deriving from currently unculturable phyla.

**Figure 1:**
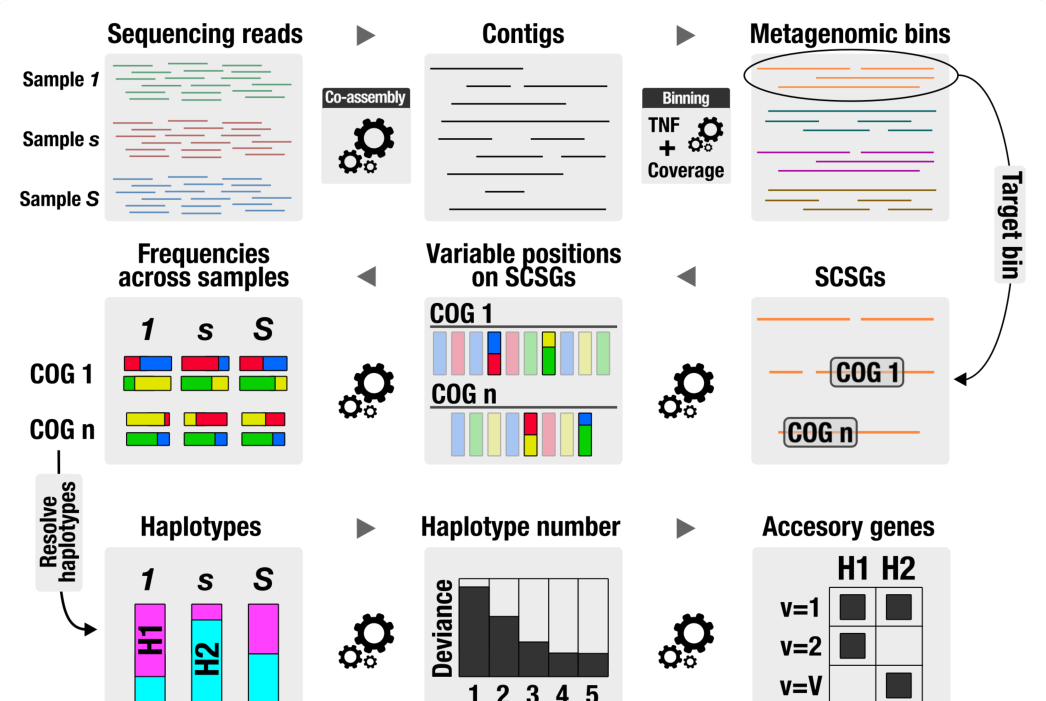
Summary of DESMAN pipeline. A full description of the statistics and bioinformatics underlying DESMAN is given in the Methods below. The software itself is available open source from https://github.com/chrisquince/DESMAN.

## Results

The DESMAN pipeline is summarised in Figure 1.

## Synthetic ‘strain’ mock

### Contig binning with CONCOCT

The assembly statistics for the synthetic ‘strain’ mock are given in Supplementary Table 2. See Supplementary Table 1 for details of the 20 genomes present. Only contigs greater than one kilobase pairs (1 kbp) in length were used for downstream analyses, and those greater than 20 kbp in length were fragmented into pieces smaller than 10 kbp [3]. CONCOCT clustered the resulting 7,545 contig fragments from these 20 genomes into 19 bins. Supplementary Figure 5 compares CONCOCT bins for each contig with the genome from which they originated. `CONCOCT` fails to separate the *E. coli* contigs that were unique to each strain correctly. However, even if the separation was perfect, contig binning could not reconstruct complete genomes for each strain, since each contig is assigned only to a single cluster. This clustering combined shared contigs across E. coli strains into bin 6, and the remaining strain-specific contigs were contained in bin 16 (Supplementary Figure 5).

To extract strains with `DESMAN`, we first combined bins 6 and 16 to recover the ‘*E. coli* pangenome’, which contained 2,028 contigs with a total length of 5,389,019 bp. We then identified genes in this contig collection using `Prodigal` [17], and assigned COGs by matching translated genes against the NCBI COG database with `RPS-BLAST`. This analysis identified 2,854 COGs, 372 of which matched with our 982 single-copy core species genes (SCSGs) for *E. coli* (see ‘Identifying core genes in target species’). These 372 SCSGs had a total length of 255,753 bp, and we confirmed that each of them occurred as single-copy in our contig collection for the *E. coli* pangenome.

### Variant detection

We mapped reads from each sample onto the contig sequences associated with the 372 SCSGs to obtain sample specific base frequencies at each position. We identified variant positions using the likelihood ratio test defined below (Equation 2) classifying positions as variants if they had a false discovery rate of less than 10^−^3. As an example, Supplementary Figure 6 displays the likelihood ratio test values for a single COG (COG0015 - Adenylosuccinate lyase) across nucleotide positions, along with true variants as determined from the known genome sequences. Supplementary Table 3 reports the confusion matrix comparing the 6,044 predicted variant positions across all 372 SCSGs with the known variants. Our test correctly recalled 97.9% of the true variant positions with a precision of 99.9% (Supplementary Table 3). Our analysis missed 125 variant positions, but manual inspection revealed that this is almost entirely due to incorrect mapping rather than the variant discovery algorithm *per se*.

### Strain deconvolution

Having identified 6,044 potential variant positions on the 372 SCSGs, we then ran the haplotype deconvolution algorithm with increasing numbers of strains *G* from 3 to 8. We ran the Gibbs sampler on 1,000 positions chosen at random with five replicate runs for each *G*, each run comprised 100 iterations of ‘burn-in’ followed by 100 samples as discussed below. The runs were initialised using the NTF algorithm with different random initialisations. At the end of the run we assigned the base sequences for each haplotype, across all positions, by generating 100 samples following 100 samples of ‘burn-in’, from these base assignments using strain frequencies and error rates from the posterior samples generated from the 1,000 positions. Figure 2A gives the posterior mean deviance, a proxy for model fit, as a function of *G*. We can see from this that the deviance decreases rapidly until *G* = 5, after which the curve flattens. In this case we can easily identify that the number of strains is indeed the five *E. coli* strains present in our mock community. We can now assess how well we can reconstruct the known sequences for *G* = 5. Supplementary Table 4 reports the number of positions at which a given haplotype differs from each reference genome in the mock data. These results confirm that each haplotype maps onto a distinct genome with error frequencies varying from 10 to 39 positions out of 6,044, representing error rates from 0.17% to 0.64% of SNP positions. The percentage of correctly predicted variable positions averaged over haplotypes was 99.58%.

**Figure 2:**
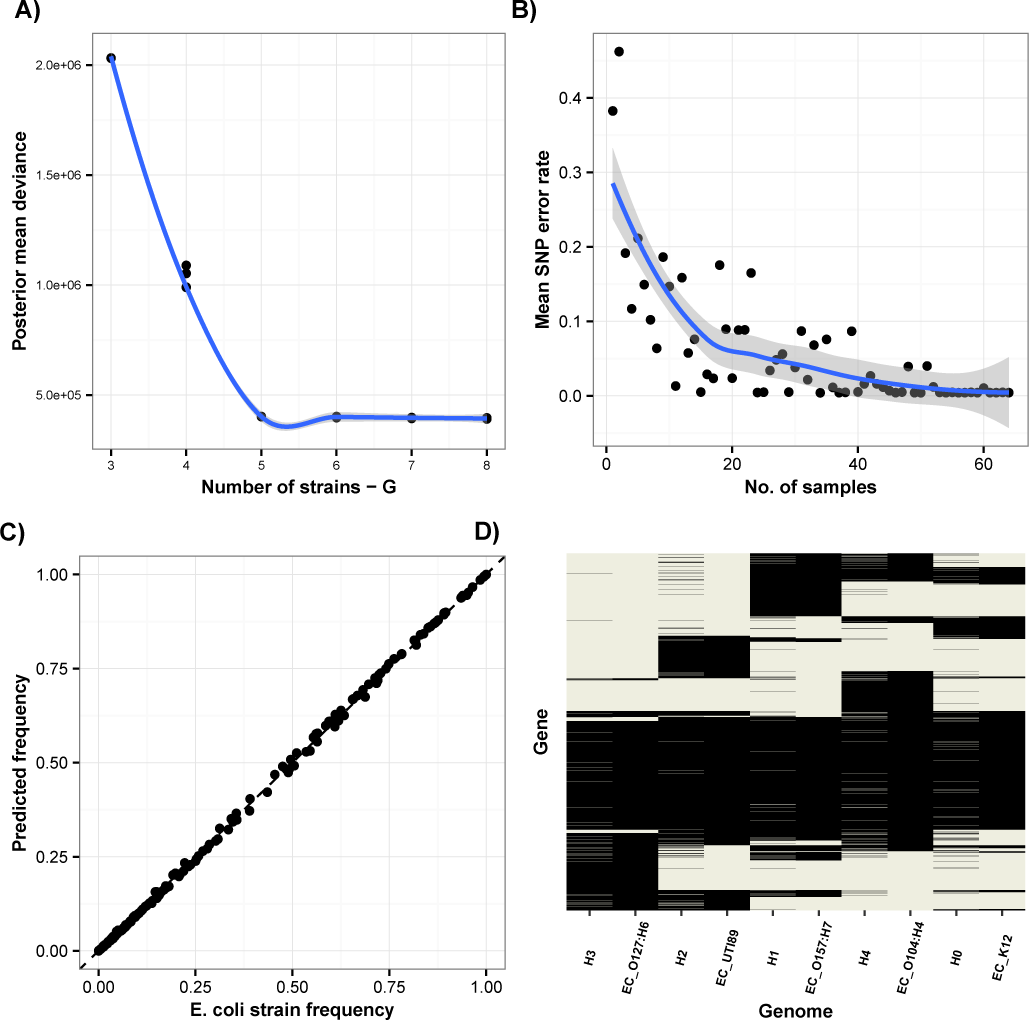
A) Posterior mean deviance for different strain number, *G*, for the synthetic ‘strain’ mock *E. coli* SCSG positions. We ran five replicates of the Gibbs sampler at each value of *G* on 1,000 random positions from the 6,044 variants identified. B) SNP accuracy as a function of sample number. The number of incorrectly inferred SNPs averaged across all five strains, and twenty replicates, of a random subset of the 64 samples. C) Comparison of true *E. coli* strain frequency vs. DESMAN predictions. We compare the known *E. coli* strain frequencies as relative coverage against the frequencies in each sample of the DESMAN predicted haplotype it mapped onto (*R*^2^ = 0.9998 and a p-value *<* 2.2e-16). D) Comparison of gene presence inferred for the haplotypes and the known assignment of genes to strain genomes. Gene presence/absence was inferred for the haplotypes using Equation 8 and compared to known references. Overall accuracy was 95.7%. These results were for the run with *G* = 5 that had the lowest posterior mean deviance.

### Comparison to existing algorithms

We also ran the `Lineage` algorithm from O’Brien *et al.* [30] on the same mock data. The model was run on the same 1,000 variants selected at random from the 6,044 variant positions we identified. We could not run the full 6,044 variant positions because of run time limitations. Their model also correctly predicted five haplotypes, however two of these were identical, and matched exactly to the EC K12 strain. Of the other three predictions, one was only 7 SNPs different from EC O104, yet the other two did not correspond to any of the true genomes. The average accuracy of prediction (the percentage of correctly predicted variable positions mapping each predicted haplotype onto the closest unique reference) was 76.32%. Supplementary Table 5 compares the `Lineage` predictions to the known strains. To provide a completely transparent comparison with `DESMAN` we also compare the `DESMAN` predictions to the known strains on just these 1,000 variant positions in Supplementary Table 6. That gave an average accuracy of 99.6%. We were unable to run `ConStrains` [27] on the same dataset, as the program complained that *insufficient coverage* of *E. coli* specific genes were obtained from the MetaPhlAn mapping. This is despite the fact that the *E. coli* coverage across our samples ranged between 37.88 and 432.00, with a median coverage of 244.00, well above the minimum of 10.0 stated to be necessary to run the `ConStrains` algorithm [27].

### Phylogenetic tree reconstruction

In Supplementary Figure 7 we display the phylogenetic analysis of 62 reference *E. coli* genomes together with the inferred strain sequences constructed using 372 SCSGs. In four out of five cases the closest relative to each strain on the tree was the genome actually used to construct the synthetic ‘strain’ mock. In the one case where it was not, *E. coli* K12, the strain was most closely related to three highly similar K12 strains including that used in the synthetic community, demonstrating that the algorithm is accurate enough to resolve strain-level phylogenetic relationships.

### Effect of sample number on strain inference

To quantify the number of samples necessary for accurate strain inference for each sample number between 1 and 64 we chose a random subset of samples that had mean strain relative abundances as similar as possible to those in the complete 64. We then ran `DESMAN` as above but only using these samples. This was done after the variant detection so all positions identified as variants were potentially included in the subsets. We ran 20 replicates of the Gibbs sampler at each sample number and then calculated SNP error rate for these runs *i.e.* the fraction of positions at which the inferred SNP differed to the true SNP in the closest matching reference. This was averaged over all five strains and 20 replicates. The results are shown together with the original 64 samples in Figure 2B, accuracy starts to decline when sample number is reduced below about 30, however, reasonable average accuracies are still achieved even with just ten samples. In addition, at low sample number accuracy is very variable across strains, typically some of the strains are resolved accurately, and others are missed completely.

### Inference of strain abundances

`DESMAN` also predicts the frequencies of each strain in each sample. We validated these predictions by comparing with the known frequencies of the *E. coli* genome each inferred strain mapped onto (Supplementary Table 4). The relative frequencies predicted by `DESMAN` are the proportion of coverage deriving from each strain. For the synthetic mock we specified the relative genome frequency of each strain in each sample, therefore we had to normalise these by the inverse of the strain genome lengths, and re-normalise. Through this analysis we obtained an almost exact correspondence between the relative frequencies for all five strains in all 64 samples (see Figure 2C). A linear regression of actual values against predictions forced through the origin resulted in a coefficient of 0.996, an adjusted *R* ^2^ = 0.9998, and a p-value *<* 2.2e-16.

### Run times

To run `DESMAN` for one choice of strain number,*G* = 5 took on average 116.86 minutes, on the synthetic ‘strain’ mock this was using ten cores on an Intel(R) Xeon(R) CPU E7-8850 v2 @ 2.30GHz. There is no parallelisation of the Gibbs sampler at the heart of `DESMAN` but replicate MCMC runs and different strain numbers can be trivially parallelised. Run time scales approximately linearly with sample number (see Supplementary Figure 8).

### Gene assignment

To validate the method for non-core gene assignment to strains in `DESMAN` we took the posterior mean strain frequencies across samples and error matrix from the run with *G* = 5 that had the lowest posterior mean deviance. These were then used as parameters to infer the presence or absence of each gene in each strain, given their mean gene coverages and frequencies of variant positions across samples (Equation 9). Figure 2D compares these inferences with the known values for each reference genome. We can determine whether a gene is present in a strain genome with an overall accuracy of 94.9%.

### *E. coli* O104:H4 outbreak

#### Assembly, contig binning, core gene identification and variation detection

The results on the synthetic ‘mock’ community are encouraging, they demonstrate that in principle `DESMAN` should be able to accurately resolve strains *de novo* from mixed populations but it can never be guaranteed that performance on synthetic data will be reproduced in the real world. There are always additional sources of noise that cannot be accounted for in simulations. Therefore, for a further test of the algorithm we applied it to 53 human fecal samples from the 2011 Shiga toxin-producing *E. coli* (STEC) O104:H4 outbreak. Here, we do not know the exact strains present and their proportions but we do know one of the strains, the outbreak strain itself from independent genome sequencing of cultured isolates [1]. So we can test our ability to resolve this particular strain.

In Supplementary Table 2 we give the assembly statistics for the *E. coli* O104:H4 outbreak data. We used the CONCOCT clustering results from the original analysis in Alneberg et al. (2014) as our starting point for the strain deconvolution. From the total of 297 CONCOCT bins we focused on just three, 95% of the contigs in which could be taxonomically assigned to *E. coli*. These bins were denoted as 83, 122 and 216 in the original nomenclature, and together they contained 2,574 contigs with a total length of 7,239 kbp. We identified 4,651 COGs in this contig collection, 673 of which matched with the 982 SCSGs that we identified above for *E. coli*. We expect that all core genes should have the same coverage profiles across samples. We can therefore compare the coverage of each putative SCSG against the median in that sample, on this basis we filtered a further 233 of these SCSGs, leaving 440 for the downstream analysis with a total length of 420,220 bp. This is an example of the extra noise arising in real samples. For the synthetic community this filtering strategy would remove no SCSGs (hence why it was not applied above).

We obtained sample-specific base frequencies at each position by mapping reads from each of the 53 STEC samples onto the contig sequences associated with the 440 SCSGs. In the following analysis we only used the 20 samples, in which the mean coverage of SCSGs was greater than five, as it would have been challenging to identify variants confidently in samples with less coverage. Aggregating frequencies across samples, we detected 28,435 potential variants (FDR *<* 1.0e-3) on these SCSGs, which were then used in the strain inference algorithm.

#### Strain deconvolution

Using these 20 samples we ran the strain deconvolution algorithm with increasing numbers of strains *G* from 2 to 10, similar to the analysis above, except that for these more complex samples we used 500 iterations rather than 100 for both the ‘burn-in’, and sampling phase. Supplementary Figure 9 displays the posterior mean deviance as a function of strain number, *G*, from this we deduce that eight strains is sufficient to explain the data.

#### Strain sequence validation

We selected the replicate run with eight strains that had the lowest posterior mean deviance, i.e. the best overall fit. To determine the reliability of these strain predictions, we compared them with their closest match in the replicate runs. Due to both random initialisation of the NTF and the stochastic nature of MCMC sampling, strains in replicates are not expected to be identical. However, the consistent emergence of similar strains across replicates increases our confidence in their prediction. The left-hand side of Figure 3 displays the comparison of each strain in the selected run to their closest match in the alternate runs, as the proportion of all SNPs that are identical averaged over positions and all four alternate replicates. This is given on the y-axis against mean relative abundance across all samples on the x-axis. From this we see that the strains fall into two groups, four relatively low abundance strains, with high SNP uncertainties *>* 20% (H1, H3, H4 and H6) and four of varying abundance that we are very confident in, each with uncertainties *<* 1% (HO, H2, H5, and H7). Results that are confirmed by the right-hand side of Figure 3, where we present a phylogenetic tree constructed from these SCSGs for the eight inferred strains and 62 reference *E. coli* genomes. For example, Strain H3 forms a long terminal branch suggesting that it does not represent a real *E. coli* strain. Similarly, H1, H4 and H6 are not nested within reference strains whereas, in contrast, the four strains with low SNP uncertainties are placed adjacent to known *E. coli* genomes. In Table Supplementary Table 7 we give the closest matching reference sequence for each strain together with nucleotide substitution rates calculated from this tree. Strain H7 is 99.8% identical to an O104:H4 outbreak strain sequenced in 2011, H5 is closely related (99.8%) to a clade mostly composed of uropathogenic *E. coli* (UPEC), and in fact all four strains that we are confident in are within 1% of a reference whereas none of the other four are. In Supplementary Figure 10 we give the relative frequencies for each of the eight inferred strains across the twenty samples with sufficient *E. coli* core genome coverage (*>* 5.0) for strain inference. Here, we have ordered samples associated with STEC by the number of days since the diarrheal symptoms first appeared. This variable is marginally negatively associated with the abundance of Strain H7, which makes sense given our identification from the core sequence that it is the 2011 O104:H4 outbreak strain.

**Figure 3:**
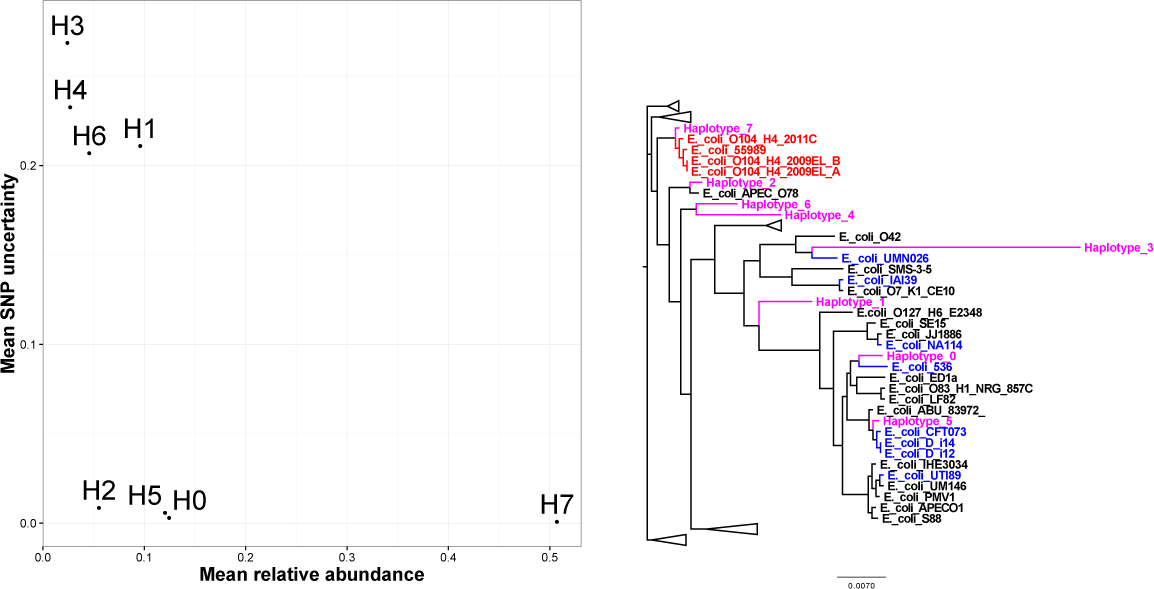
Validation of reconstructed strains for the *E. coli* O104:H4 outbreak. Left) The ‘mean SNP uncertainty’,*i.e.* the proportion of SNPs that a strain differs from its closest match in a replicate run, averaged over all the other replicates. This is shown on the y-axis against mean relative abundance across samples on the x-axis. Right) Phylogenetic tree constructed for the eight inferred strains found for the *E. coli* O104:H4 outbreak. The SCSGs for the strains and reference genomes were aligned separately using `mafft`[19], trimmed and then concatenated together. The tree was constructed using `FastTree`[33]. Inferred strains are shown as magenta, O104:H4 strains in red and uropathogenic *E. coli* (UPEC) in blue. Both results were for the run with *G* = 8 that had the lowest posterior mean deviance.

#### Anaerobic digestion reactor metagenomes

The analysis of the *E. coli* O104:H4 outbreak proves that on real data `DESMAN` is capable of reconstructing valid strains, even when sample number is relatively small. It is potentially very useful to be able to resolve strains direct from epidemic metagenome data but typically for common human pathogens we have extensive strain databases that could be used in a supervised ap-proach instead. The real appeal of `DESMAN` is its ability to extract strain variation for the uncultured microbes that represent the majority of diversity in environmental samples. To demonstrate this we applied the pipeline to a total of 95 metagenome samples taken from three replicate laboratory anaerobic digestion (AD) bioreactors converting distillery waste into biogas (*Quince et al. Unpublished data*). The focus here is not to consider the engineering implications of this study but simply to demonstrate our ability to resolve the diversity present at high resolution in this complex microbial community.

#### Assembly, contig binning, core gene identification and variation detection

The 95 reactor samples were sequenced on two flow cells of a HiSeq 2500 generating a total of 521,492,655 2*×* 125 bp reads. They were assembled with Ray using a kmer size of 41, the assembly statistics are given in Supplementary Table 2. Contigs greater than 2*kbp* in length were split if they exceeded 20*kbp* as described in [3]. All 186,081 resulting contig fragments were clustered by `CONCOCT` [3]. A total of 355 bins were generated by this process.

Genes were called on these contigs and annotated to COGs as above [10]. Through the analysis of 36 single copy core COGs (SCGs), previously identified to be found in all bacterial genomes in a single copy [3], we determined that 139 of these bins contained bacterial genomes that had greater than 75% of these genes present and in a single copy. That is they were at least 75% pure and complete. These we identified as metagenome assembled genomes (MAGs) and used them in the downstream analysis. The SCG frequencies for all bins are shown in Supplementary Figure 12.

In resolving strain diversity we restricted our analysis to 26 AD MAGs with a total coverage across reactor biomass samples of greater than 75.0. Less total coverage than this and it is unlikely that the strain inference would work accurately. These MAGs are summarised in Supplementary Table 8, they represent a broad range of diversity deriving from nine different bacterial and archaeal phyla, the most frequently represented being the bacterial Proteobacteria with 6 representatives, followed by the archaeal Euryarchaeota with 5. However, much less well studied phyla are also present including 4 Chloroflexi, and 2 Spirochaetes and even one MAG from the currently unculturable Candidate Phyla Radiation (CPR) which we tentatively assign to Wolfebacteria [7].

For these environmental organisms, we have no reference genomes available so we run `DESMAN` using just the 36 SCGs that we can be confident are core and single copy in almost every microbe. As is evident from Supplementary Table 8, some MAGs will miss some SCGs but this is not critical it just reduces the number of positions that we can perform inference on. The number of SCGs used for each MAG were further reduced by filtering those with outlying coverages, the numbers of SCGs before and after filtering and total length in bp are given in Supplementary Table 9 for each MAG. The median number of SCGs after filtering was 18.5. We then ran variant detection on these filtered SCGs, the frequency of variants varied considered between MAGs ranging from 0.4% to 9.6% with a median of 2.75% indicating a substantial degree of genetic variation within the organisms. This variation did not correlate with genome size but did show a weak positive relationship with number of KEGG metabolic modules encoded in the MAG (see Supplementary Figure 13 -*R* ^2^ = 0.1132,*p − value* = 0.05). This is in contrast to the suggestion based on just two assembled genomes that metabolic complexity should be associated with reduced genetic variability [28]. Most of this association is driven by the fact the two MAGs with very simple metabolisms the CPR MAG and one assigned to Firmicutes had very low levels of variation.

#### Strain deconvolution

Some of the variation in these organisms will represent genetic variation within a population, but some will be attributable to the presence of multiple strains [30]. We inferred strains for each of these MAGs, running five replicates of the Gibbs sampler for from 1 to 5 strains. The best fitting number of strains was determined by inspection of the deviance plots as above. Then we only considered a strain as valid if it had a mean relative abundance of greater than 5% across samples and a SNP uncertainty less than 5%. In total 16 out of the 26 MAGs had multiple strains,*i.e.* 61.5%, in most cases two strains were inferred, and in the two cases when we predicted 4 strains, we could only be confident of 2 (see Supplementary Table 10). Therefore in all cases we are comparing the predominant strain with a single variant in each MAG. We can calculate the divergence between the two strains in terms of percentage nucleotide difference on the core genes, this varies between 0.159% and 4.258%, with a median of 1.051%.

#### Comparison of nucleotide divergence and genome divergence

A key question to ask is how does divergence in core gene nucleotide identity relate to similarity in the accessory genome between the strains for each of the 16 pairs. To address this we inferred for each strain pair both gene presence/absence for all genes in the MAG and the sequences of the genes present in each strain. We then clustered genes at 5% nucleotide identity and calculated the percentage of gene clusters that differed between the strains. The comparison of this with core nucleotide divergence is shown in the top panel of Figure 4. There is no clear relationship between these two quantities (linear regression,*R* ^2^ = 0.048, *p − value* = 0.2064). Since for any given species we would expect the two to be positively correlated this result implies that between species the degree of correlation must vary. We can validate this by examining environmental species for which multiple strains are available, in Supplementary Figure 14 we show nucleotide divergence against genome divergence for three environmental organisms (*Methanosarcina mazei*,*Lactococcus lactis* and *Acinetobacter pittii*). This confirms that the results in Figure 4 are reasonable and that whilst core nucleotide divergence and genome divergence do correlate, there is a great deal of variation within species, and the relationship between the two varies from one species to another. In particular, in *Acinetobacter pittii* more genome divergence is observed for the same level of nucleotide divergence than for the other two.

**Figure 4:**
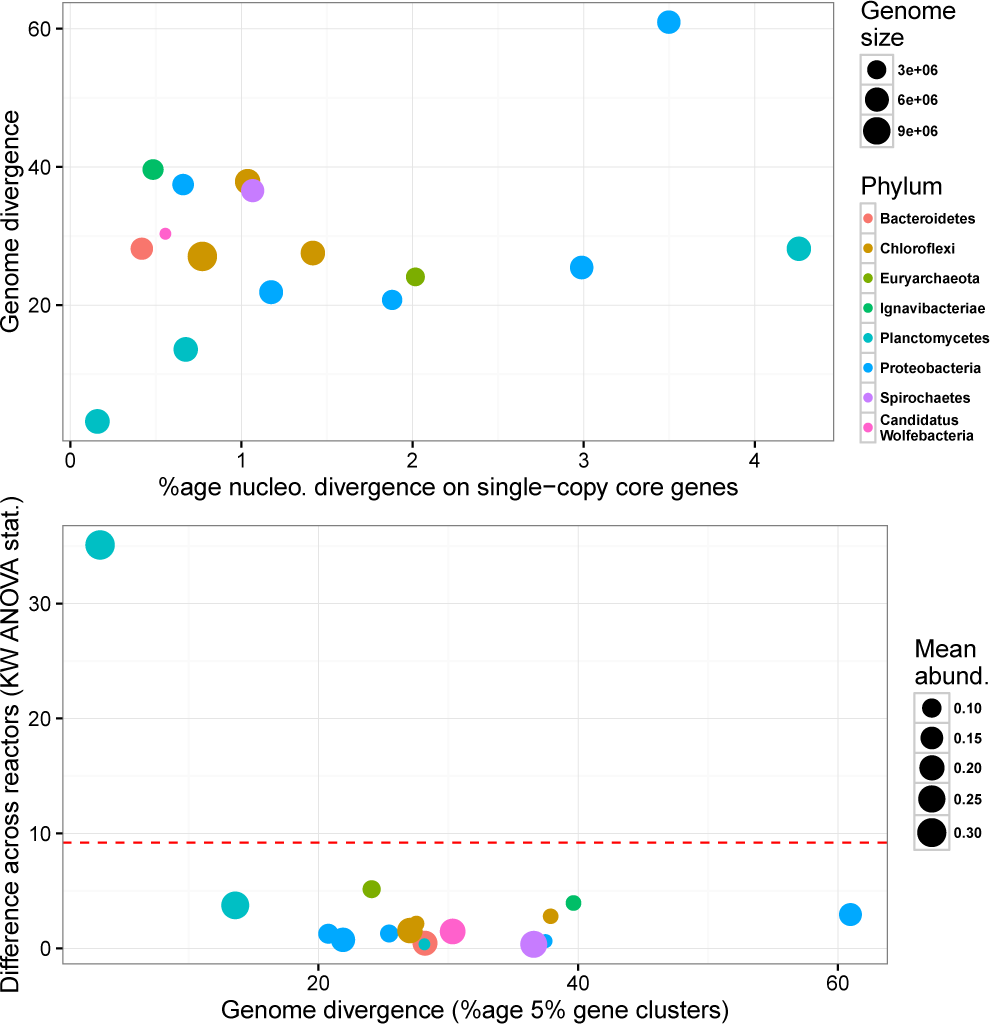
Top). Comparison of nucleotide divergence on core genes with 5% gene cluster divergence for 16 strain pairs. For each pair of strains we calculate percentage nucleotide divergence on the core genes and compare to percentage of 5% gene clusters differing between the two. Bottom) Comparison of 5% gene cluster divergence with difference in strain abundances across replicate reactors. For each strain pair we used Kruskal-Wallis ANOVA to test for a significant difference in minor strain abundance between replicate reactors, the chi-squared test-statistic is shown on the y-axis against genome divergence, the red dashed line corresponds to significance at *p* = 0.05.

#### Strain niche partitioning

For most strain resolved MAGs, there is no significant difference in the minor strain abundance between the three replicate reactors. In the bottom panel of Figure 4 we plot the chi-squared test-statistic for the null hypothesis of no difference in strain abundance across reactors against genome divergence between the strains. In fact, only one MAG, Cluster158 from the phylum Planctomycetes, exhibits strains that differ significantly in abundance between the reactors (see Supplementary Figure 15 - left panel). This suggests that the strains in the other 15 MAGs are not ecologically redundant to each other, otherwise their relative proportions would differ between reactors due to stochastic effects in inoculation, and fluctuate in abundance over time. In fact their abundances are either constant over time or change deterministically with a consistent trend (e.g. Supplementary Figure 15 right panel). What is striking about Cluster 158, is that its strains have an almost identical gene complement with a genome divergence of just 3.171%, the next lowest divergence is 13.59%. This is very suggestive and may indicate that ecological redundancy in environmental strains is only possible when there is very little difference in gene complement. Otherwise different strains have different roles reflected by differing accessory genes and stable or deterministically varying proportions due to niche partitioning.

## Discussion

We have demonstrated on both *in silico* and real data sets the ability of DESMAN to correctly infer and reconstruct microbial strains from metagenomic data *de novo* by using subtle nucleotide variations in mapping results. Besides resolving strains that were initially binned together, our approach also reported their relative abundances in samples, elucidating biologically relevant patterns, such as evidence for niche partitioning in the anaerobic digestion reactors amongst strains that had a sufficiently divergent gene complement. `DESMAN` can resolve strains at any level of divergence, and can link fragmented sequences.

`DESMAN` was more effective at reconstructing the five strains of *E. coli* in our mock dataset than `Lineage` [30]. This is surprising, as the model used in `Lineage` aims to exploit an additional level of information that is not used in our algorithm through the simultaneous construction of a phylogenetic tree between strains. This, in theory, should enable a more powerful inference of haplotype dissimilarities across sites, but the average SNP accuracy of the `Lineage` predicted haplotypes was just 76.32% compared to 99.58% accuracy for `DESMAN`. We have introduced a novel method based on non-negative tensor factorisation (NTF) for initialising our inference algorithm. Since MCMC sampling can be sensitive to initial conditions, it is possible that this methodological improvement provides an advantage over the `Lineage` model. Alternatively, the improvement we achieved could be due to `DESMAN’s` use of a more complex error model, or the fully conjugate Gibbs sampler. Nevertheless, it would be worthwhile to extend `DESMAN` to include phylogenetic information, or conversely, introduce some of our improvements into the `Lineage` algorithm. This would further improve our collective ability to resolve complex pangenomes *de novo* from metagenomic assemblies. We were unable to run the `ConStrains` algorithm on our data, which in itself illustrates the advantage of a strategy, in which we separate the steps of mapping, variant calling and haplotype inference. Although we suspect the partially heuristic and non-probabilistic approach utilised in `ConStrains` would have been unable to compete with the fully Bayesian algorithm employed in `DESMAN`.

The underlying haplotype inference model in `DESMAN` could be improved. Position-dependent error rates may be relevant given that particular sequence motifs are associated with high error rates on Illumina sequencers [34]. More fundamentally, we could develop models that do not assume independence across variant positions combining information from the cooccurrence of variants in the same read with the modelling of strain abundances across multiple samples. This could be particularly relevant as single molecule long read sequencers such as Nanopore become more commonly used [26]. In addition, it would have been preferable to have a more principled method for determining the number of strains present, rather than just examining the posterior mean deviance. This could be achieved through Bayesian non-parametrics such as a Dirichlet process prior for the strain frequencies allowing a potentially infinite number of strains to be present, with only a finite but flexible number actually observed [29]. Alternatively a variational Bayesian approach could be utilised to obtain a lower bound on the marginal likelihood and this used to distinguish between models [11].

To the best of our knowledge, this is the first study to demonstrate that coverage across multiple samples can be used to infer contig counts across strains within a pangenome. This critical step enables the identification and recovery of true genomic diversity present in metagenomic data. This allowed us to resolve strain diversity and gene complement in entirely uncultured species including members of the Candidate Phyla Radiations [7].

The `DESMAN` pipeline is an open source software, and is available via the URL https://github.com/chrisquince/DESMAN.

## Models and Methods

The `DESMAN` (De novo Extraction of Strains from MetAgeNomes) pipeline is a strategy for resolving both strain haplotypes and variations in gene content directly from short-read shotgun metagenome data. Our proposed approach comprises commonly employed steps of an assembly-based metagenomic binning workflow (such as co-assembly of the data, annotation of resulting contigs, mapping short reads to the assembly, and identification of genome bins), followed by preparing genome bins that match to the target organism for strain extraction using the novel `DESMAN` algorithm described below.

## Assembly and mapping

The first step is to co-assemble all reads from all samples. As chimeric contigs can confound the downstream analyses with DESMAN, the choice of assembler and the assembly parameters are important to target more accurate contigs rather than longer, but potentially chimeric ones, even if these selections result in relatively lower N50 values for the overall assembly. For our analyses we used idba ud [31] or Ray [6]. The result of an assembly will be a set of *D* contigs with lengths in base pairs *L_d_*, and sequence composition *U_d_* with elements *u_d,l_* drawn from the set of nucleotides *{A, C, G, T}*.

Following co-assembly we used `bwa mem` [24] to map raw reads in each sample individually back onto the assembled contigs. We then used `samtools` [23], and `sequenza-utils` [14] to generate a 4-dimensional tensor *𝒩* reporting the observed base frequencies, *n_d,l,s,a_*, for each contig and base position in each sample, where *d* = 1,…, *D*, *l* = 1,…, *L_d_*, *s* = 1,…, *S*, and *a* = 1,…, 4 that represents an alphabetical ordering of bases 1 → *A*, 2 → *C*, 3 → *G*, 4 → *T*.

Using this tensor we calculated an additional *D* × *S* matrix, giving the mean coverage of each contig in each sample as:

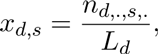

where we have used the convenient ‘dot’ notation for summation, i.e. 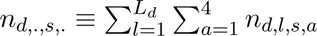

## Contig clustering, and target species identification

To identify putative species for strain extraction *de novo*, we recommend grouping contigs into bins using a clustering algorithm that takes both sequence composition, and differential coverage of contigs into consideration, for this task we used CONCOCT [3]. Here, we assume one or more of the resulting bins match to the target species, and contain a total of *C* contigs with indices that are a subset of {1,…, *D*}. For convenience, here we re-index the coverages and base frequency tensor such that *x_c,s_*, and *n_c,l,s,a_*, gives the mean coverage, and base frequencies in this subset, respectively.

## Identifying core genes in target species

The algorithm assumes a fixed number of strains in the target species. However, in general, not every gene in every contig will be present in all strains. We address this by identifying a subset of the sequences that occur in every strain as a single copy. Here we identify those ‘core genes’ for *Escherichia coli* by (1) downloading 62 complete *E. coli* genomes from the NCBI, and (2) assigning Clusters of Orthologous Groups of proteins (COGs) [10] to the genes in these genomes. This allowed us to identify 982 COGs that are both single-copy, and had an average of greater than 95% nucleotide identity between the 62*E. coli* genomes genomes. We denote these COGs as single-copy core species genes (SCSGs). We then searched for SCSGs in MAGs that represent our target species, and created a subset of the variant tensor with base positions that fall within SCSGs hits. We denote this subset as *n_h,l,s,a_*, where *h* is now indexed over the *H* SCSGs found and *l* is the position within each SCSG from 1,…, *L_h_*, which have lengths *L_h_*. We denote the coverages of these genes as *x_h,s_*.

For thec *E. coli* analyses we have reference genomes available, this is not the case for the anaerobic digester (AD) metagenomes, there we used a completely *de novo* approach, using 36 single-copy core genes (SCGs) that are conserved across all species [3] but any other single-copy gene collection [8, 12] could serve for the same purpose. The result is a decrease in resolution due to the decreased length of sequence that variants are called on but it is still sufficient to resolve strains at low nucloetide divergence.

In real data sets, we have noticed that some core genes will in some samples have higher coverages than expected. We suspect that this is due to the recruitment of reads from low abundance relatives that fail to be assembled. To account for this we apply an additional filtering step to the core genes. All core genes should have the same coverage profile across samples. Therefore we applied a robust filtering strategy based around the median absolute deviation [22]. We calculated the absolute divergence of each gene coverage from the median denoted 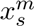:

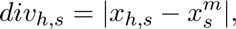

and then the median of these divergences, denote that 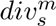. If:

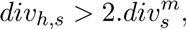

we flag it as an outlier in that sample. We only use genes that are not flagged in at least a fraction *f* of samples, where in these analyses *f* was set at 80%.

## Variant detection

Our algorithmic strategy begins with a rigorous method for identifying possible variant positions within the SCSGs. The main principle is to use a likelihood ratio test to distinguish between two hypotheses for each position. The null hypothesis *ℋ*_0_ is that the observed bases are generated from a single true base under a multinomial distribution and an error matrix that is position-independent. We denote this error matrix *∊* with elements *∊_a,b_* giving the probability that a base *b* is observed when the true base is *a*. The alternative hypothesis *ℋ*_1_ in which *ℋ*_0_ is nested is that two true bases are present. For the purposes of this test we ignore the distribution of variants over samples, working with the total frequency of each base across all samples:

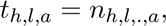

although the generalisation of our approach to multiple samples would be quite straightforward, we chose not to do this for computational reasons, and because we achieve sufficient variant detection accuracy aggregating frequencies.

If we make the reasonable assumption that *∊_a,a_* > *∊_a,b_* for *b* ≠ *a* for all *a*, then for a single true base with errors, the maximum likelihood solution for the true base is the consensus at that location which we denote by the vector *M_h_* for each SCSG with elements:

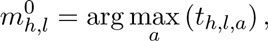

the likelihood for *ℋ*_0_ at each position is then the multinomial assuming that bases are independently generated under the error model:

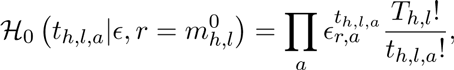

where we use 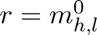 to index the maximum likelihood true base. Similarly, for the two bases hypothesis, the maximum likelihood solution for the second base (or variant) is:

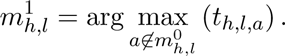

Then the likelihood for the hypothesis *ℋ*_1_ at each position is:

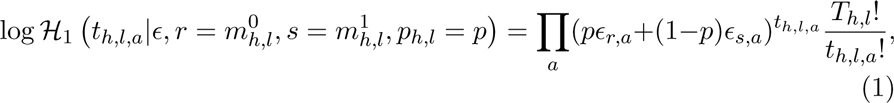

where we have introduced a new parameter for the relative frequency of the consensus base,*p*, and *T_h,l_* is the total number of bases at the focal position, *T_h,l_* = *t_h,l,._*. We set a minimum lower bound on this frequency corresponding to the minimum observable variant frequency, *p_l_* < *p* ≤ 1, for the synthetic ‘mock’ community we set *p_l_* = 0.01 *i.e.* 1%, for the other two real data sets where we want to be more conservative we used *p_l_* = 0.03. For each position we determine this by maximum likelihood performing a simple one dimensional optimisation of Equation 1 with respect to *p*. Having defined these likelihoods, our ratio test is:

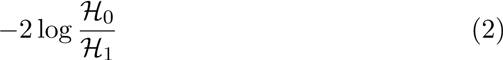

which will be approximately distributed as a chi-squared distribution with one degree of freedom. Hence, we can use this test to determine p-values for the hypothesis that a variant is present at a particular position.

There still remains the question of how to determine the error matrix, *∊*. We assume that these errors are position independent, and adopt an EM like iterative approach to its determination. We start with a rough approximation to *∊*, categorise positions as variants or not, and then recalculate *∊* as simply the observed base transition frequency across all non-variant positions. We then re-classify positions and repeat until *∊* and the number of variants detected converge. Finally, we apply Benjamini-Hochberg correction to account for multiple testing to give a false discovery rate (FDR) or q-value for a variant at each position [4].

## Probabilistic model for variant frequencies

Having identified a subset of positions that are likely variants the next step of the pipeline is to use the frequencies of those variants across multiple samples to link the variants into haplotypes. We use a fairly low q-value cut-off for variant detection, using all those with a false FDR < 1.0*e* − 3. This ensures that we limit the positions used in this computationally costly next step to those most likely to be true variants. The cost is that we may miss some low frequency haplotypes but these are unlikely to be confidently determined anyway. We will index the variant positions on the SCSGs by *v* and for convenience keep the same index across SCSGs which we order by their COG number, so that v runs from 1,…, *N*_1_,…, *N*_1_ + *N*_2_,…, ∑_*h*_ *N_h_* where *N_h_* is the number of variants on the hth SCSG and keep a note of the mapping back to the original position and SCSG denoted *υ* → (*l_υ_, h_υ_*). We denote the total number of variants by *V* = ∑_*h*_ *N_h_* and the tensor of variant frequencies obtained by subsetting *n_h_υ_,l_υ_,s,a_* → *n_υ,s,a_* on the variant positions as *N*.

## Model likelihood

The central assumption behind the model is that these variant frequencies can be generated from *G* underlying haplotypes with relative frequencies in each sample *s* denoted by *π_g,s_*, so that *π_.,s_* = 1. Each haplotype then has a defined base at each variant position denoted *τ_υ,g,a_*. To encode the bases we use 4-dimensional vectors with elements ∈ {0, 1} where a 1 indicates the base and all other entries are 0. The mapping to bases is irrelevant but we use the same alphabetical ordering as above, thus *τ_υ,g,._* = 1.

We also assume a position-independent base transition, or error matrix giving the probability of observing a base *b* given a true base *a* as above, *∊_a,b_*. Then, assuming independence across variant positions, i.e. explicitly ignoring any read linkage, and more reasonably between samples, the model likelihood is simply a product of multinomials:

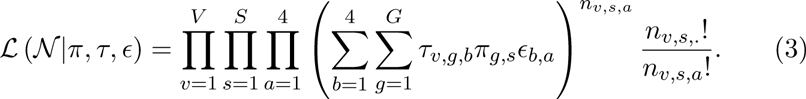

## Model priors

Having defined the likelihood, here we specify some simple conjugate priors for the model parameters. For the frequencies in each sample we assume symmetric Dirichlet priors with parameter:

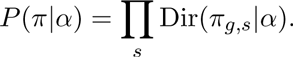

Similarly, for each row of the base transition matrix we assume independent Dirichlets:

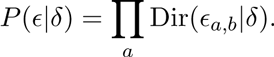

with parameter *δ*. Finally, for the haplotypes themselves (*τ*), we assume independence across positions and haplotypes, with uniform priors over the 4 states:

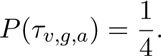

## Gibbs sampling strategy

We will adopt a Bayesian approach to inference of the model parameters generating samples from the joint posterior distribution:

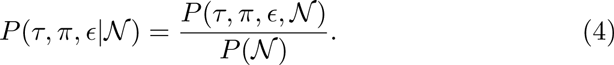

We use a Gibbs sampling algorithm to sample from the conditional posterior of each parameter in turn, which will converge on the joint posterior given sufficient iterations [5]. The following three steps define one iteration of the Gibbs sampler:

1. The conditional posterior distribution for the haplotypes,*τ_v,g,a_*, is just:

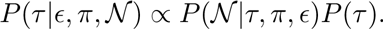 Each variant position contributes independently to this term, so we can sample each position independently. The haplotype assignments are discrete states so their conditional will also be a discrete distribution. We sample *τ* for each genome in turn, from the conditional distribution for that genome, with the assignments of the other genomes fixed to their current values:

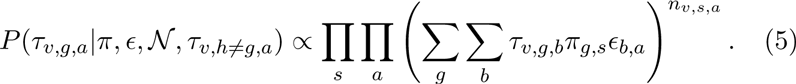
2. To sample the *∊* we introduce an auxiliary variable, *ν,_s,a,b_*, which gives the number of bases of type *a* that were generated by a base of type *b* at location *υ* in sample *s*. Its distribution conditional on *τ*,*π*,*∊* and *𝒩* will be multinomial:

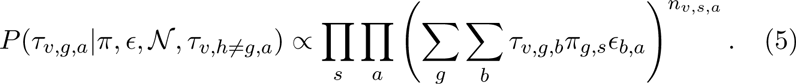

where:

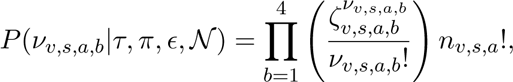 Since the multinomial is conjugate to the Dirichlet prior assumed for *∊* then we can easily sample *∊* conditional on *ν*:

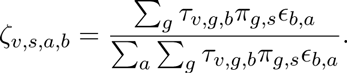
3. To sample *π* we define a second auxiliary variable *ξ_υ,s,a,b,g_* which gives the number of bases of type *a* that were generated by a base of type *b* at each position *υ* from haplotype *g* in sample *s*. This variable conditioned on *τ*,*π*,*∊* and will be distributed as:

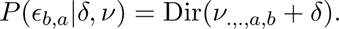

with:

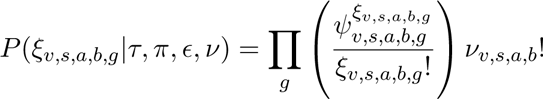 Similarly, *π* is also Dirichlet distributed conditional on *ξ*:

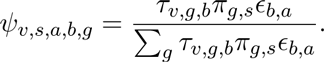

## Initialisation of the Gibbs sampler

Gibbs samplers can be sensitive to initial conditions. To ensure rapid convergence on a region of high posterior probability, we consider a simplified version of the problem. We calculate the proportions of each variant at each position in each sample:

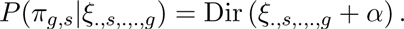

Then an approximate solution for *τ* and *π* will minimise the difference between these observations, and:

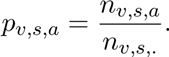

If we relax the demand that *τ_v,g,a_* ∈ 0, 1, and instead allow it to be continuous, then solving this problem is an example of non-negative tensor factorisation (NTF), which itself is a generalisation of the better known non-negative matrix factorisation problem (NMF) [39]. We adapted the standard multiplicative update NTF algorithm that minimises the generalised Kullback-Leibler divergence between *p* and 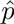:

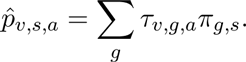

This is equivalent to assuming that the observed proportions are a sum of independent Poisson distributed components from each haplotype, ignoring the issue that the Poisson is a discrete distribution [9]. The standard multiplicative NMF algorithm can be applied to our problem [21] by rearranging the *τ* tensor as a 4*V × G* matrix τ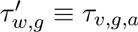, where *w* =*υ* + (*a* − 1)*V*. By doing so, we have created a matrix from the tensor by stacking each of the base components of all the haplotypes vertically. Similarly, we rearrange the variant tensor into a 4*V × S* matrix with elements 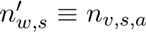, where *w* = *υ* + (*a* − 1)*V*. The update algorithms become:

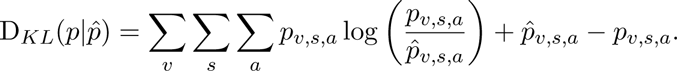

Then we simply add a normalisation step:

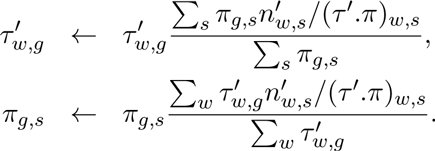

Having run the NTF until the reduction in D_*KL*_ was smaller than 10^−5^, we discretised the predicted *τ* values such that the predicted base at each position for each haplotype was the one with the largest *τ*^′^. We used these values with the *π* as the starting point for the Gibbs sampler.

## Implementation of the Gibbs sampler

In practice, following initialisation with the NTF, we run the Gibbs sampling algorithm twice for a fixed number of iterations, the first run is a ‘burn-in’ phase to ensure convergence, which is checked via manual inspection of the time series of parameter values. The second run is the actual sampler, from which *T* samples, are stored as samples from the posterior distribution,*θ_t_* =(*τ_t_, π_t_, ∊_t_*) with *t* = 1,…, *T*. These can then be summarised by the posterior means, 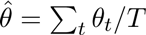, and used in subsequent downstream analysis. We also store the sample with the maximum log-posterior, denoted *θ*^*^ = (*τ*^*^, *π*^*^, ε^*^),if a single most probable sample is required. For many data sets *V* will be too large for samples to be generated within reasonable time. Fortunately, we do not need to use all variant positions to calculate *π* with sufficient accuracy. We randomly selected a subset of the variants, ran the sampler, obtained samples (*π_t_*, *∊_t_*), and use these to assign haplotypes to all positions, by running the Gibbs sampler just updating *τ* sequentially using Equation 5 and iterating through the stored (*π_t_, ∊_t_*).

## Determining the number of haplotypes and haplotype validation

Ideally the ‘Bayes factor’ or the model evidence, the denominator in Equation 4, would be used to compare between models with different numbers of haplotypes. Unfortunately, there is no simple reliable procedure to accurately determine the Bayes factor from Gibbs sampling output. For this reason we suggest examining the posterior mean deviance [15]:

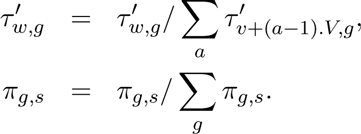

as the number of haplotypes increases the model will fit better and *D* will decrease, when the rate of decrease is sufficiently small then we conclude that we have determined the major abundant haplotypes or strains present. This method is ambiguous but has the virtue of not making any unwarranted assumptions necessary for approximate estimation of the Bayes factor. To validate individual haplotypes we compare replicate runs of the model. Since the model is stochastic then different sets of haplotypes will be generated each time. If in replicate runs we observe the same haplotypes then we can be confident in their validity. Therefore calculating the closest matching haplotypes across replicates gives an estimate of our confidence in them. We define the ‘mean SNP uncertainty’ for a haplotype as the fraction of positions for which it differs from its closest match in a replicate run, averaged over all the other replicates.

## Resolving the Accessory Genome

Having resolved the number of strains and their haplotypes on the core genome, we now consider the question of how to determine the accessory genome for each strain. The strategy below could equally well be applied to either contigs or genes called on those contigs. In our experience, contigs are frequently chimeric, and we have achieved better results with gene based approaches. If contig asssignments are required then a simple consensus of the genes on a contig can be used. We will therefore describe a gene based analysis keeping in mind that contigs could be used interchangeably.

We should have already assigned genes on all contigs in the target bin or bins above. Now we consider not just the SCSGs but all genes which we will index *f* = 1*,…, F*. Just as for the SCSGs we can identify variant positions on the total gene set using Equation 2. In fact, we apply a slightly modified version of this strategy in this case because of the large number of positions to be screened, replacing the 1D optimisation of *p* with an estimation of the frequency of the consensus base as the ratio of the observed number of consensus bases to the total, 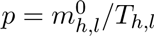.

We will denote the number of variant positions associated with gene *f* by 𝒩*_f_*. In this case we do need to keep track of which variant maps to which gene explicitly so we will consider a 4D variant tensor denoted *𝓜* with elements *m_f,l,s,a_* where *l* is indexed from 1,…, *𝒩_f_*. This is generated by simply subsetting the original contig variant tensor *𝒩*, to the variants associated with each gene. In practice, to speed up the algorithm we only use a random subset of variants (20 was used here), since all variants contain the information necessary to determine which gene is present in which strain. An additional source of information we will use is the average coverage of each gene across samples, this is the exact analogue of the contig coverage introduced above and we will denote it with the same symbol,*i.e.* χ with elements *x_f,s_*.

Determining the accessory genome corresponds to inferring the copy number of each gene in each strain. We denote this integer as *η_f,g_*, for each of the genes *f* = 1,…, *F* associated with the species in each strain, *g* = 1,…, *G*. The ideas we present here could be extended to multi-copy genes, however, the current implementation of DESMAN assumes that all genes are present in zero or one copies, *η_f,g_* ∈ {0, 1}. This simplifies the implementation considerably and in real assemblies the vast majority of genes are either present or absent in a strain, for example for the STEC genome this is true of 98.8% of the genes.

The first step is to determine the likelihood, we assume that this is separable for the variants and the coverages. This is an approximation as the variant positions will contribute to the mean coverage calculation. Formally we assume:

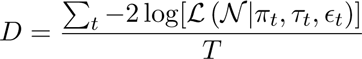

The first likelihood is as Equation 3 a product of multinomials:

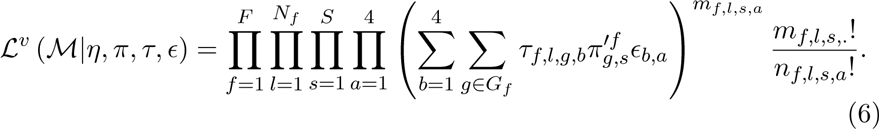

The difference is that now the sum over the strains *g* are only those for which *η_f,g_* > 0, those which actually possess a copy of gene *f*, a set which we denote *g* ∈ *G_f_*. The relative frequencies then have to be renormalised accordingly so that:

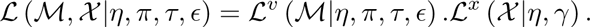

The likelihood for the coverages is somewhat simpler. We know the relative proportions of each strain in each sample, *π_g,s_*, we also know the mean total coverage on the core genes:

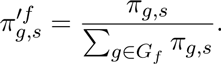

Therefore, we can calculate the coverage associated with each strain:

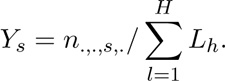

We can make the approximation that each copy of a contig from a strain contributes independently to the total mean coverage observed for that contig in a particular sample. If we further assume that this contribution is Poisson distributed with mean *γ_g,s_*, then the total contribution will be from the superposition property of Poisson distributions, again Poisson with mean *λ_f,s_* = ∑_*g*_ *η_f,g_γ_g,s_*. Thus:

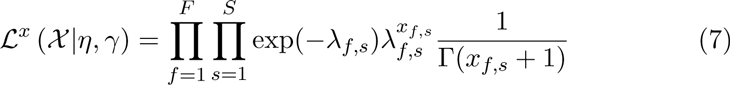

Our strategy to sample the gene assignments *η_f,g_* is to keep the the relative proportions of each strain in each sample, *π_g,s_*, and the error matrix, *∊_b,a_* fixed at their posterior mean values 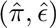. We then use a Gibbs sampling strategy to jointly sample both the *η_f,g_* and the haplotypes of those strains *τ_f,l,g,a_*. In general, we assume a geometric prior for the *η_f,g_* so that 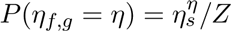, where *η_s_* is less than one to penalise multi-copy genes,although here as mentioned above we restrict ourselves to binary *η*, and *Z* is a normalisation constant. Each gene contributes to the likelihood independently and so can be sampled independently, we can therefore loop through the genes, sampling the *η* for each strain conditional on the other genomes fixed at their current values:

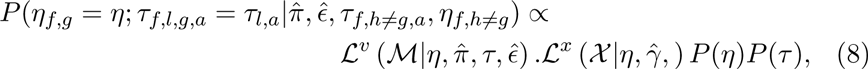

substituting Equations 6 and 7 into this and using uniform priors for *τ*.

In order to improve the speed of convergence of this sampler we developed an approximate strategy to initialise *η_f,g_* using just the coverages, *x_f,s_*. If we ignore for now that *η_f,g_* is discrete, then the maximum likelihood prediction for *η_f,g_* from Equation 7 will correspond to minimising the generalised Kullback-Leilber divergence between the observed coverages *x_f,s_*, andtheir predictions, 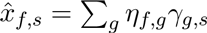:

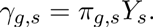

This also corresponds to non-negative matrix factorisation but with a fixed estimate for *γ_g,s_*. Therefore, to solve it for *η_f,g_*, we only need one of the multiplicative update rules [21]:

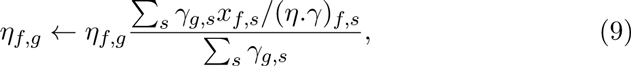

which gives continuous estimates for *η_f,g_*, but we simply round these to the nearest integer for discrete copy number predictions.

The sampler is initialised using Equation 9 before applying a ‘burnin’ and sampling phase using Equation 8. Typically we have found that a relatively small number of samples, just 20, are sufficient before the *η* values converge. We also only use a random subset of the variant positions (again 20) for the sampling as discussed above. Optionally we then allow an additional sampling phase to determine the remaining *τ*, the haplotype sequences, with the fixed at their posterior mean values, if required.

## Author’s contributions

CQ wrote the DESMAN code, developed statistics, performed analyses and wrote the first draft of the MS. SC, SG, and GC designed and performed the AD reactor experiments. SR helped develop the Gibbs sampler. JA helped develop the analysis pipeline. ME provided the original motivation for the algorithm, contributed to the codebase, and generated figures. All authors read and approved the final manuscript.

## Acknowledgements

CQ is funded through an MRC fellowship (MR/M50161X/1) as part of the Cloud Infrastructure for Microbial Bioinformatics (CLIMB) consortium (MR/L015080/1). SC, SGS and GC, and the bioreactor trials, were supported by the Engineering and Physical Sciences Research Council, UK (EP/J00538X/1 - A Global Solution to Transforming Waste). GC was supported by a European Research Council Starting Grant (3C-BIOTECH 261330). AME was supported by Frank R. Lillie Research Innovation Award. Béatrice Laroche provided useful comments that improved the Gibbs sampler.

---

**Table S1.**
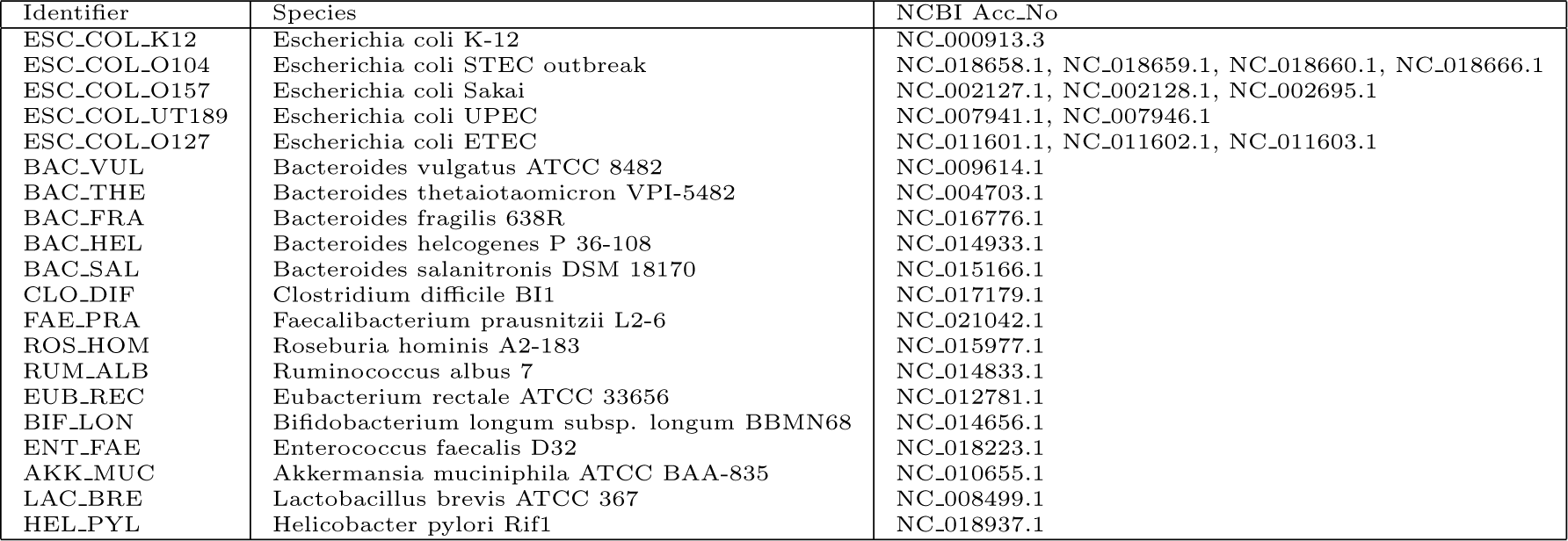
The 20 genomes used in the strain sythetic mock community.

**Table S2:** Co-assembly statistics for the three data sets used in this study. The *E. coli* O104:H4 outbreak and AD reactor data sets were assembled using Ray with kmer size 41 [6]. The synthetic ‘strain’ mock was assembled using idba ud with default parameters and kmer sizes increasing from 21 to 121 and the ‘pre correction’ flag. Contigs greater than 20kbp in length were fragmented into 10kbp pieces. The N50 length was calculated for contigs greater than 1kbp.

| Data set | No. samples | No. reads | Total length | No. contigs > 1kbp | No. contigs > 2kbp | N50 length |
| --- | --- | --- | --- | --- | --- | --- |
| ‘Strain’ mock | 64 | $1.504 \times 10^9$ | 60Mbp | 7,545 | 6,002 | 10,000 |
| *E. coli* O104:H4 outbreak | 53 | $2.2 \times 10^8$ | 317 Mbp | 142,723 | 47,918 | 2,402 |
| AD reactors | 95 | $5.21 \times 10^8$ | 1.394 Gbp | 417,253 | 186,081 | 5,810 |

**Table S3.** Confusion matrix for variant detection in the ‘strain’ mock. A contingency table or confusion matrix giving the frequency of positions that were (rows - True) or were not (rows - False) predicted to be variants with a FDR or q-value *<* 1.0e-3 that actually were (columns - True) or were not (columns - False). Overall prediction accuracy was 99.9% with a precision of 99.9% and a recall of 97.9%.

|  | Actual |  |
| --- | --- | --- |
| Predicted | False | True |
| False | 249,584 | 125 |
| True | 6 | 6,038 |

**Table S4:** Comparison of inferred strains to true reference genomes in the ‘strain’ mock. This table compares the predicted strains for one run of the algorithm with the five known *E. coli* genomes at each of the 6,044 variant positions. The integer frequencies give the number of positions where each strain (row) differs from a genome (column). The overall accuracy rate for correct prediction of SNPs at each position averaged over strains was 99.58%.

|  | Strain genomes |  |  |  |  |
| --- | --- | --- | --- | --- | --- |
| Predicted | EC_O127 | EC_O104 | EC_O157 | EC_K12 | EC_UT189 |
| H_0 | 3532 | 1606 | 2083 | 16 | 3603 |
| H_1 | 3728 | 2306 | 39 | 2084 | 3753 |
| H_2 | 1715 | 3681 | 3755 | 3587 | 35 |
| H_3 | 10 | 3637 | 3718 | 3522 | 1728 |
| H_4 | 3628 | 27 | 2288 | 1591 | 3681 |

**Table S5:** Comparison of inferred strains to true reference genomes in the ‘strain’ mock using the Lineage algorithm[30] for 1,000 variant positions. This table compares the predicted strains for one run of the Lineage algorithm with the five known *E. coli* genomes at each of 1,000 variant positions chosen at random from the 6,044 variant positions. The integer frequencies give the number of positions where each strain (row) differs from a genome (column). The overall accuracy rate for mapping of strains onto genomes was 76.32%.

|  | Strain genomes |  |  |  |  |
| --- | --- | --- | --- | --- | --- |
| Predicted | EC_O127 | EC_O104 | EC_O157 | EC_K12 | EC_UT189 |
| H_0 | 147 | 741 | 740 | 736 | 138 |
| H_1 | 600 | 243 | 333 | 0 | 631 |
| H_2 | 600 | 243 | 333 | 0 | 631 |
| H_3 | 705 | 405 | 162 | 177 | 726 |
| H_4 | 601 | 7 | 354 | 236 | 632 |

**Table S6.** Comparison of inferred strains to true reference genomes in the ‘strain’ mock using the DESMAN algorithm for 1,000 variant positions. This table compares the predicted strains for one run of the DESMAN algorithm with the five known *E. coli* genomes at each of 1,000 variant positions chosen at random from the 6,044 variant positions. The integer frequencies give the number of positions where each strain (row) differs from a genome (column). The overall SNP accuracy rate for mapping of strains onto genomes was 99.6%.

|  | Strain genomes |  |  |  |  |
| --- | --- | --- | --- | --- | --- |
| Predicted | EC_O127 | EC_O104 | EC_O157 | EC_K12 | EC_UT189 |
| H_0 | 600 | 243 | 334 | 0 | 634 |
| H_1 | 629 | 359 | 9 | 331 | 643 |
| H_2 | 277 | 635 | 646 | 631 | 5 |
| H_3 | 2 | 604 | 628 | 598 | 282 |
| H_4 | 602 | 4 | 355 | 239 | 636 |

**Table S7:** Closest matching reference genomes for the seven inferred STEC strains from *E. coli* O104:H4 outbreak. Inferred strain sequences were aligned with references and a phylogenetic tree constructed for the eight inferred strains found for the *E. coli* O104:H4 outbreak. The SCSGs for the strains and reference genomes were aligned separately using `mafft`[19], trimmed and then concatenated together. The tree was constructed using `FastTree`[33]. Here we give for each strain the closest reference in terms of phylogenetic distance,*d*, or total average nucleotide substitutions.

| Strain | Closest ref. | $d$ | 1.0 - $d$ |
| --- | --- | --- | --- |
| H_0 | *E. coli* 536 | 0.00758 | 0.99242 |
| H_1 | *E. coli* O127 H6 E2348 | 0.02117 | 0.97883 |
| H_2 | *E. coli* APEC O78 | 0.00305 | 0.99695 |
| H_3 | *E. coli* UMN026 | 0.04244 | 0.95756 |
| H_4 | *E. coli* APEC O78 | 0.02039 | 0.97961 |
| H_5 | *E. coli* ABU 83972 | 0.00195 | 0.99805 |
| H_6 | *E. coli* APEC O78 | 0.01411 | 0.98589 |
| H_7 | *E. coli* O104 H4 2011C | 0.00165 | 0.99835 |

**Table S8.**
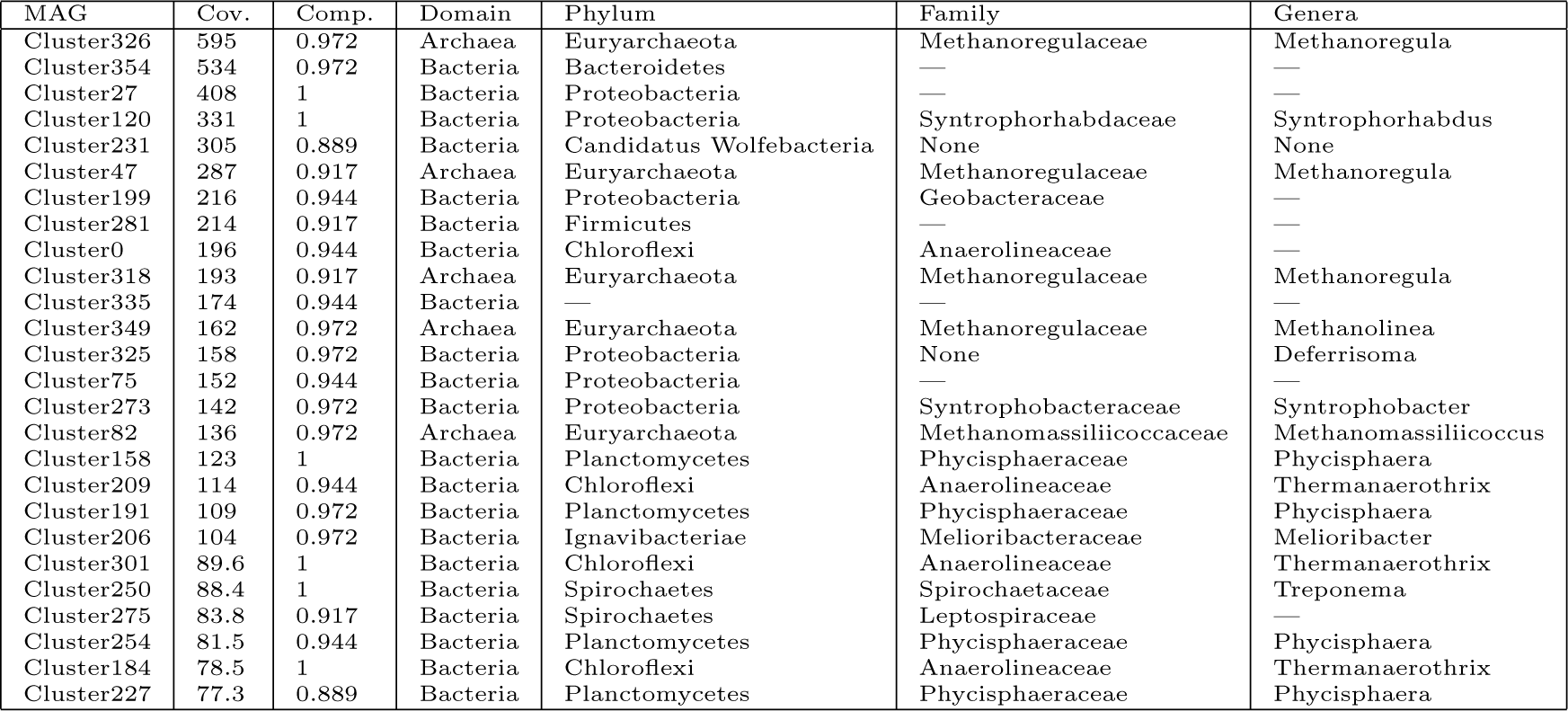
The 24 MAGs used in the AD metagenome strain analysis. These 26 metagenome assembled genomes (MAGs) were processed through the DESMAN pipeline. We give the total coverage of the MAG across the (Cov.) 43 biomass samples, the other 52 samples corresponded to influent, efluent and inocula. We also give the completeness of the MAG as the proportion of 36 single-copy core genes (SCGs) that were present in a single copy, together with taxonomic assignments based on a phylogeny constructed with those core genes, and 1755 reference genomes. A dash (—) indicates that a confident assignment could not be made at that level.

**Table S9.** Variants on single-copy core genes (SCGs) for 26 anaerobic digestion (AD) MAGs. For each of the 26 AD MAGs we give the number of single copy core genes in the MAG (no. SCGs), the number that remain after filtering based on median coverage (no. filter.), the length in bp of the filtered SCGs (len. filter.), the number of variants detected (no. var.), and the fraction of variants per bp. (f. var.).

| MAG | no. SCGs | no. filter. | len. filter. | no. var. | f. var. |
| --- | --- | --- | --- | --- | --- |
| Cluster326 | 35 | 16 | 18678 | 548 | 0.029 |
| Cluster354 | 35 | 21 | 17643 | 538 | 0.03 |
| Cluster27 | 36 | 23 | 19227 | 130 | 0.007 |
| Cluster120 | 36 | 18 | 19467 | 747 | 0.038 |
| Cluster231 | 32 | 20 | 15288 | 182 | 0.012 |
| Cluster47 | 33 | 23 | 22470 | 304 | 0.014 |
| Cluster199 | 34 | 21 | 17583 | 1280 | 0.073 |
| Cluster281 | 33 | 25 | 20253 | 146 | 0.007 |
| Cluster0 | 34 | 15 | 15171 | 1452 | 0.096 |
| Cluster318 | 33 | 21 | 19830 | 464 | 0.023 |
| Cluster335 | 34 | 17 | 14760 | 196 | 0.013 |
| Cluster349 | 35 | 20 | 20493 | 185 | 0.009 |
| Cluster325 | 35 | 19 | 18978 | 767 | 0.04 |
| Cluster75 | 34 | 17 | 16434 | 753 | 0.046 |
| Cluster273 | 35 | 21 | 20391 | 1005 | 0.049 |
| Cluster82 | 35 | 23 | 21621 | 1191 | 0.055 |
| Cluster158 | 36 | 16 | 19572 | 314 | 0.016 |
| Cluster209 | 34 | 16 | 16989 | 448 | 0.026 |
| Cluster191 | 35 | 21 | 21342 | 676 | 0.032 |
| Cluster206 | 35 | 21 | 21867 | 81 | 0.004 |
| Cluster301 | 36 | 15 | 17160 | 117 | 0.007 |
| Cluster250 | 36 | 18 | 20031 | 1810 | 0.09 |
| Cluster275 | 33 | 17 | 18291 | 673 | 0.037 |
| Cluster254 | 34 | 17 | 20196 | 928 | 0.046 |
| Cluster184 | 36 | 18 | 19353 | 265 | 0.014 |
| Cluster227 | 32 | 17 | 17622 | 293 | 0.017 |

**Table S10.**
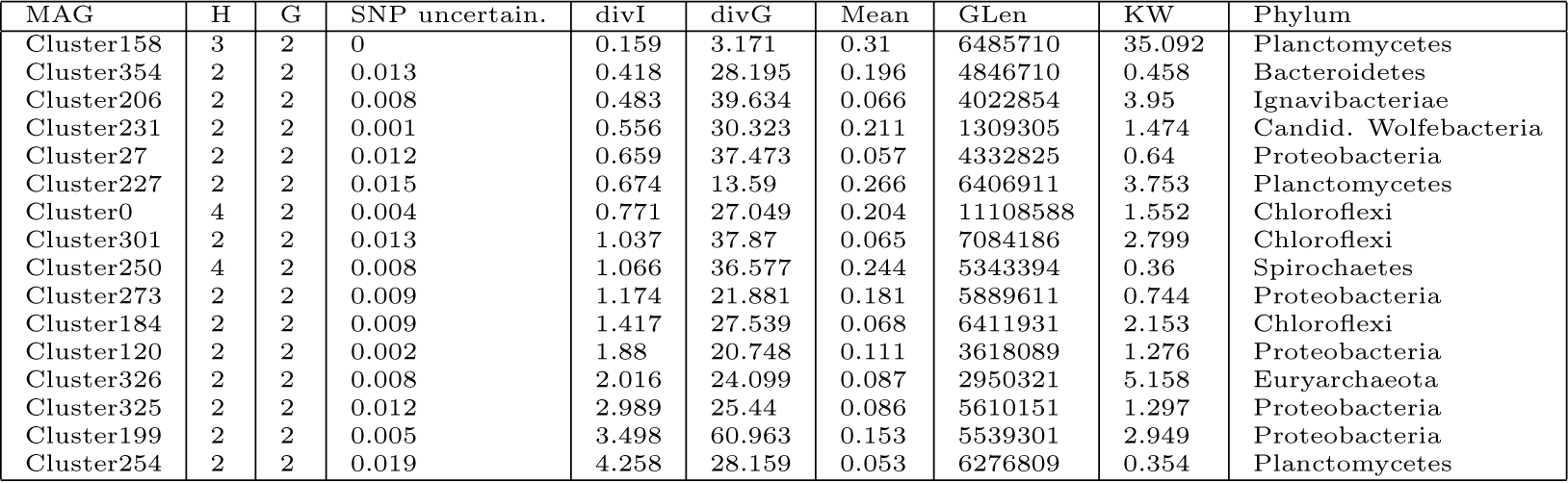
Strain inference for 16 anaerobic digestion (AD) MAGs. Here we give details of the inferred strains for the 16 of 26 MAGs for which multiple strains were observed. For each MAG,*G* is the number of strains in the best fit,*H* is the number following removal of those with less than 5% mean abundance across samples and a SNP uncertainty greater than 5% (in all cases *H* = 2). We also give the mean SNP uncertainty across strains (SNP uncertain.), the % nucleotide difference between the two strains (divI), the % difference in 5% gene overlap (divG), the mean abundance of the less abundant strain (Mean), the MAG genome length (GLen), the difference between abundances in the three replicates by Kruskal-Wallis ANOVA (KW), and assigned phyla (Phylum).

**Figure S5.**
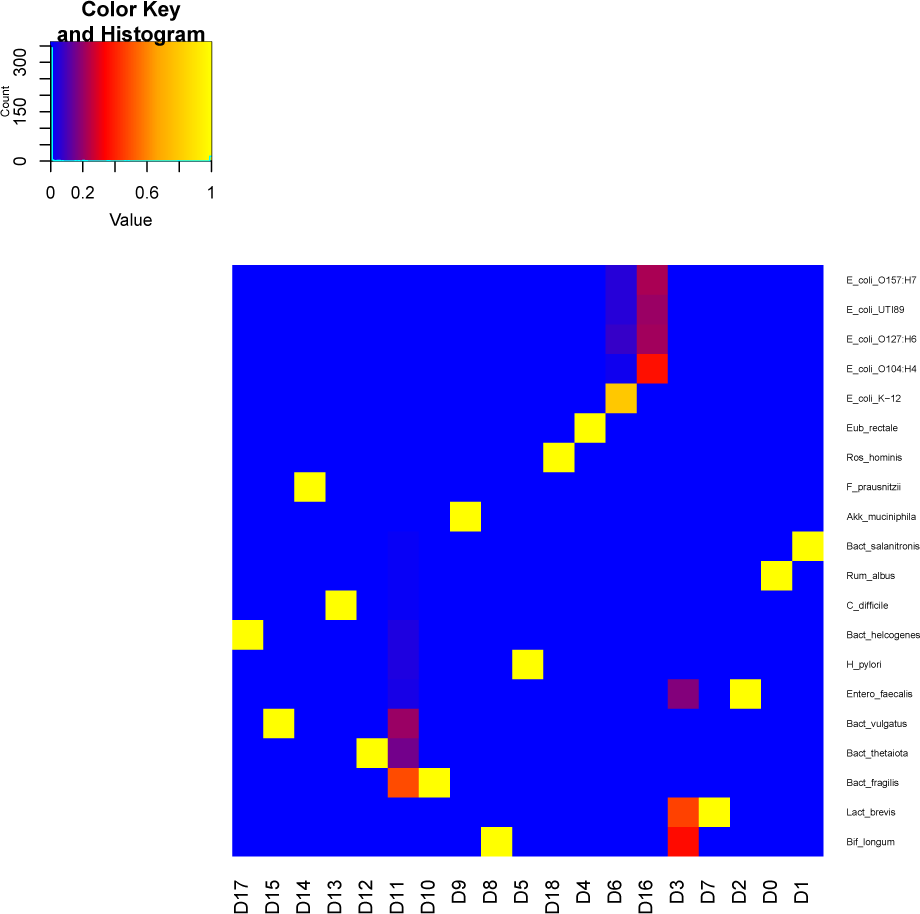
Confusion matrix for the synthetic ‘strain’ mock. Heat-map where intensity indicates proportion of contigs in each CONCOCT cluster that derive from each strain. There were 19 clusters and 20 strains. The recall was 98.1% and precision 96.1% comparing the genome labels and clusters [3] with an overall accuracy of 0.971 as given by the adjusted Rand index.

**Figure S6.**
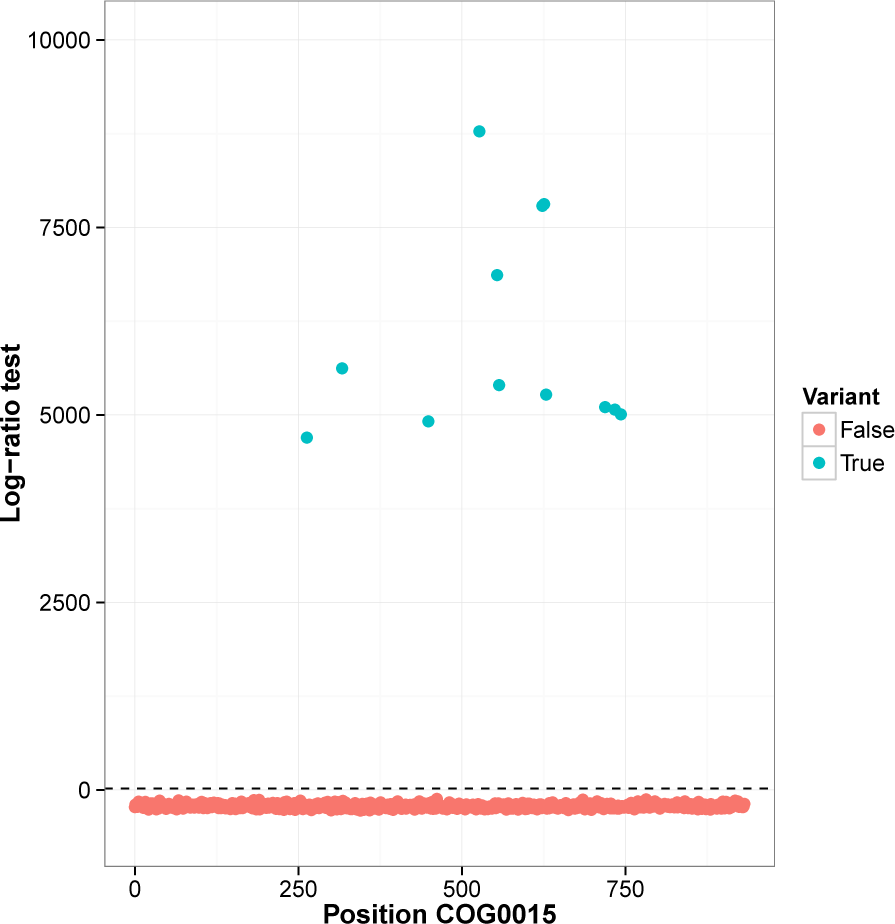
Log-ratio test statistic for variants on COG0015 of the synthetic ‘strain’ mock. The likelihood ratio test statistic (Equation 2) for positions along COG0015 - Adenylosuccinate lyase. Positions that are true variants are coloured blue, positions with no variation, red. The dashed line corresponds to a FDR or q-value of 1.0e-3. Positions above this line are classified as variants under the test. Note negative log-ratios occur because the minimum variant frequency *p_L_* is set at 1%.

**Figure S7.**
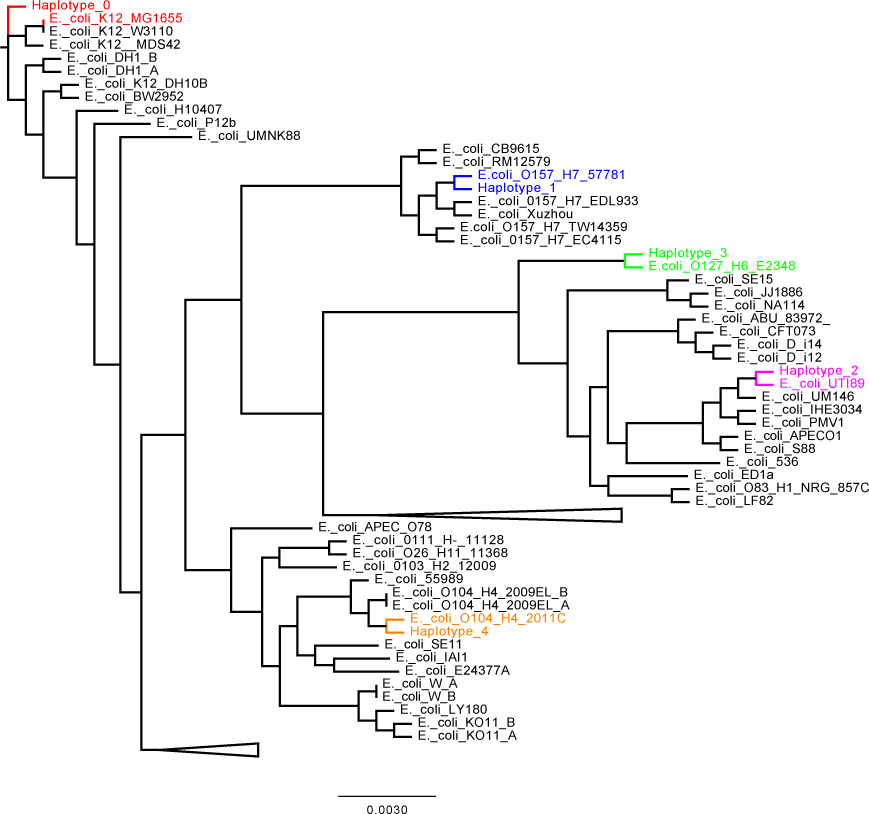
Phylogenetic tree constructed for the five inferred strains found in the ‘synthetic strain’ mock and 62*E. coli* reference sequences. The 372 SCSGs for the strains and reference genomes were aligned separately using mafft[19], trimmed and then concatenated together. The tree was constructed using FastTree[33]. The known reference genome each strain mapped onto from Supplementary Table 4 is shown in the same colour as the corresponding strain. These results were for the run with *G* = 5 that had the lowest posterior mean deviance.

**Figure S8.**
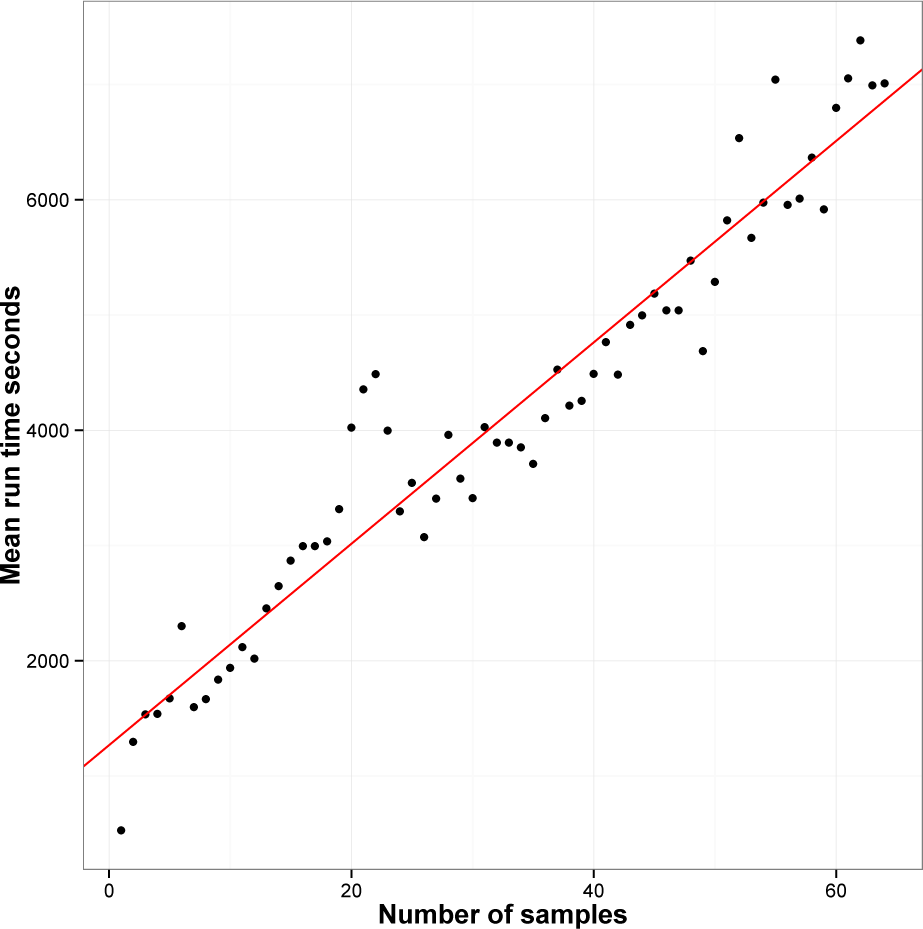
Run times for DESMAN in seconds for increasing sample number. The total run time for DESMAN on the synthetic ‘strain’ mock is shown as a function of number of samples. Results are the mean time averaged over twenty replicates comprising random subsamples of the original 64 samples. Each run was for *G* = 5 and comprised ten threads run in parallel on a Intel(R) Xeon(R) CPU E7-8850 v2 @ 2.30GHz.

**Figure S9.**
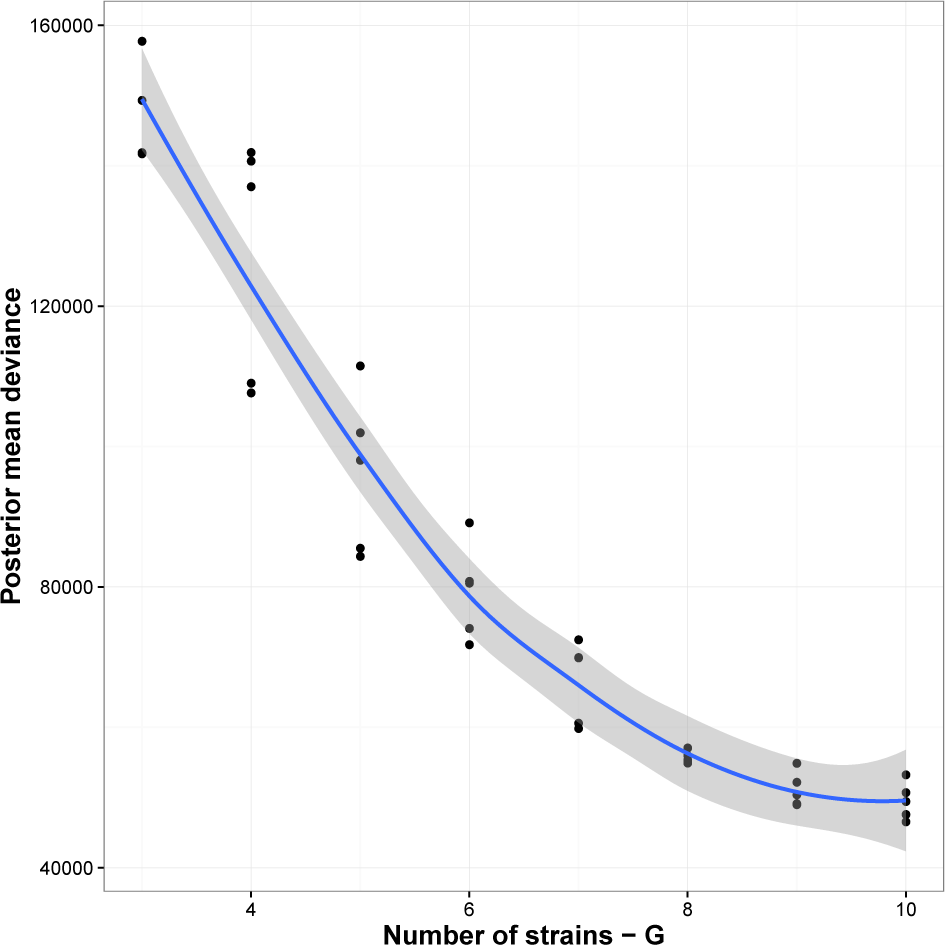
Posterior mean deviance as a function of *G* for the *E. coli* O104:H4 outbreak SCSG positions. The posterior mean deviance as a function of *G* for five replicates running the strain resolution Gibbs sampler for 1,000 random positions from the 28,435 potential variants identified on the 440 SCSGs for the *E. coli* clusters in the *E. coli* O104:H4 outbreak.

**Figure S10.**
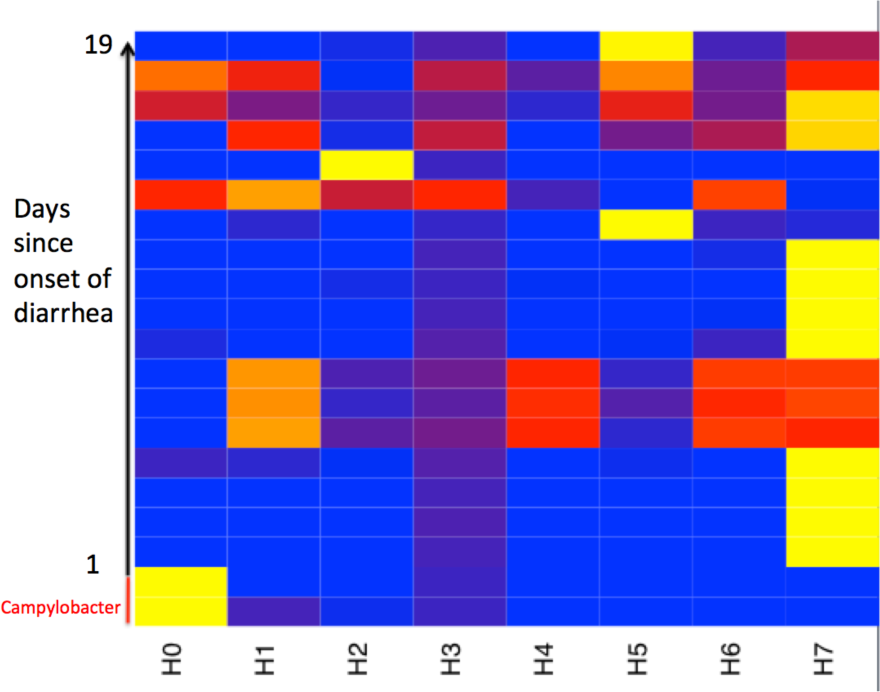
strain relative abundances across samples for the STEC *E. coli* O104:H4 outbreak. The relative frequencies of the eight predicted *E. coli* strains in the twenty samples from the STEC *E. coli* O104:H4 outbreak that had a mean coverage across *E. coli* SCSGs greater than five. The two samples at the bottom of the heat map derived from a *Campylobacter jejuni* infected individual, the other 18 samples were all determined to be infected with STEC. We have ordered these by ‘Days since onset of diarrhea (ddays)’, the number of days ago that the individual first experienced diarrhea symptoms. The relative abundance of strain 7 negatively correlates with *ddays* (*τ* =*−* 0.366,*p* = 0.0414, Kendall’s tau coefficient).

**Figure S11.**
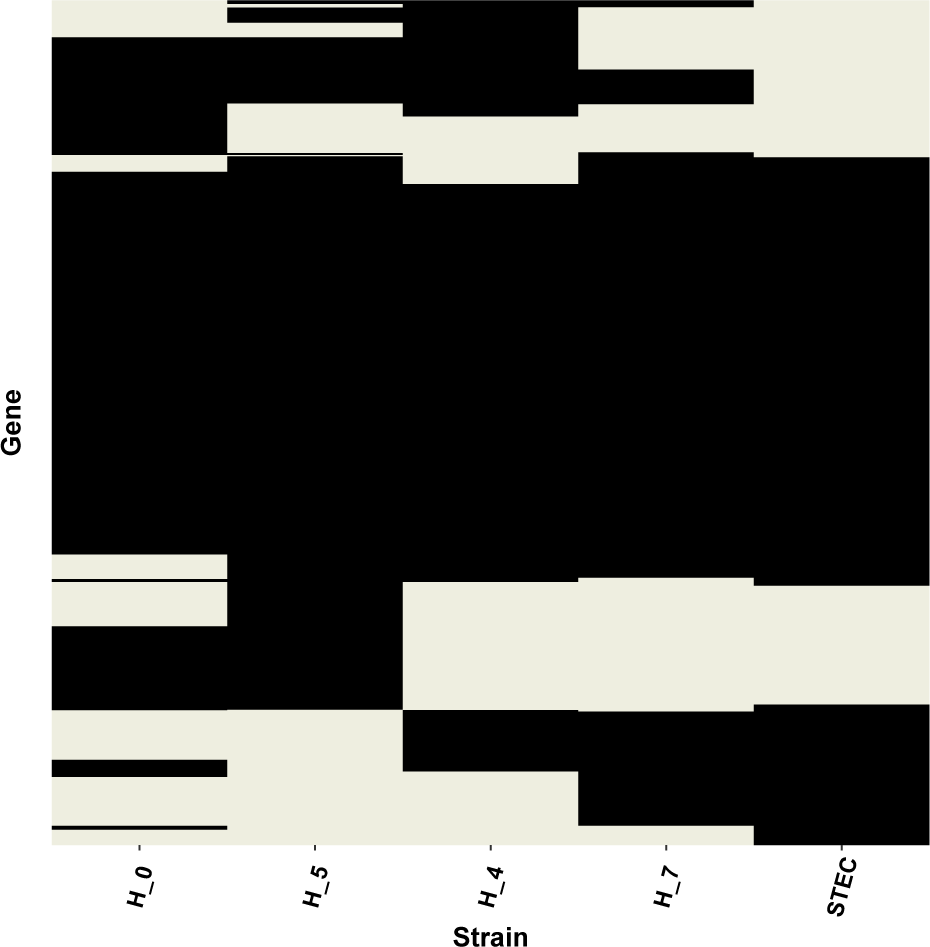
Comparison of gene presence/absence inferred for the four strains we are confident in from the STEC *E. coli* O104:H4 outbreak together with results for the known outbreak genome. Gene presence/absences were inferred for the strains using Equation 8. They were calculated for the STEC genome (*Escherichia coli* O104:H4 str. 2011C) by mapping using MUMmer. Comparing *ℋ*_7_ and the STEC genome, 91.8% of counts matched. These results were for the run with *G* = 8 that had the lowest posterior mean deviance.

**Figure S12.**
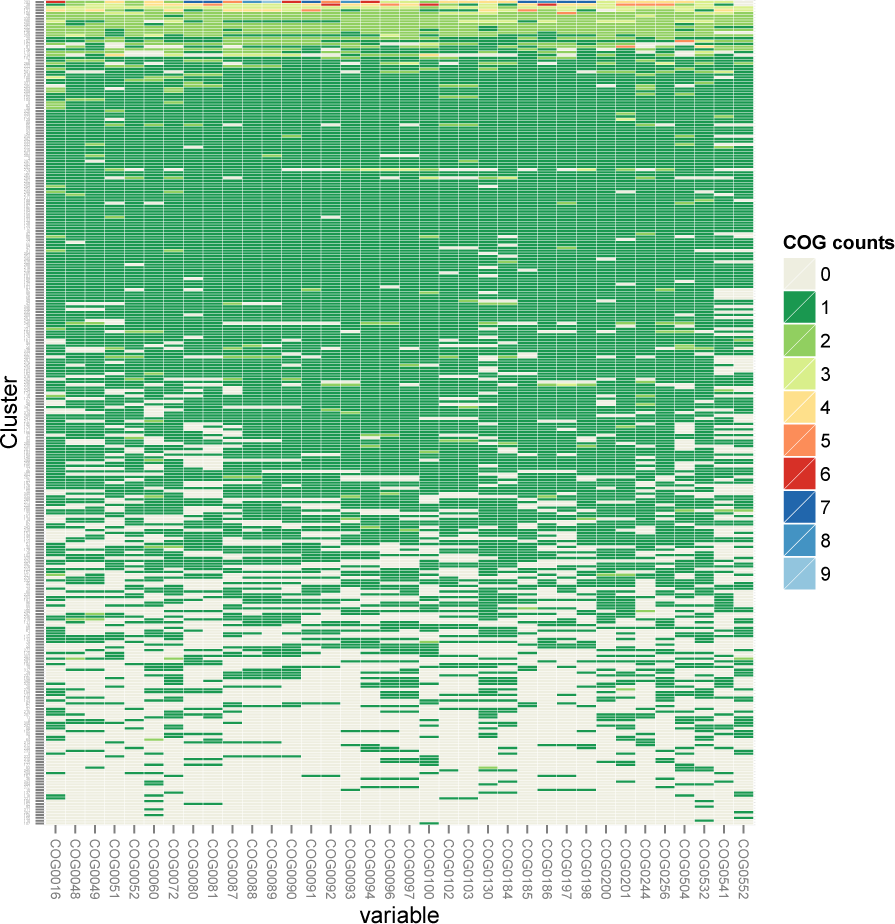
Single core gene frequencies in the 236 CONCOCT clusters generated by binning all contigs *>* 2*kbp* in length from the AD metagenome samples. 139 of these clusters were at least 75% pure and complete. Each row corresponds to a cluster and columns are SCGs.

**Figure S13.**
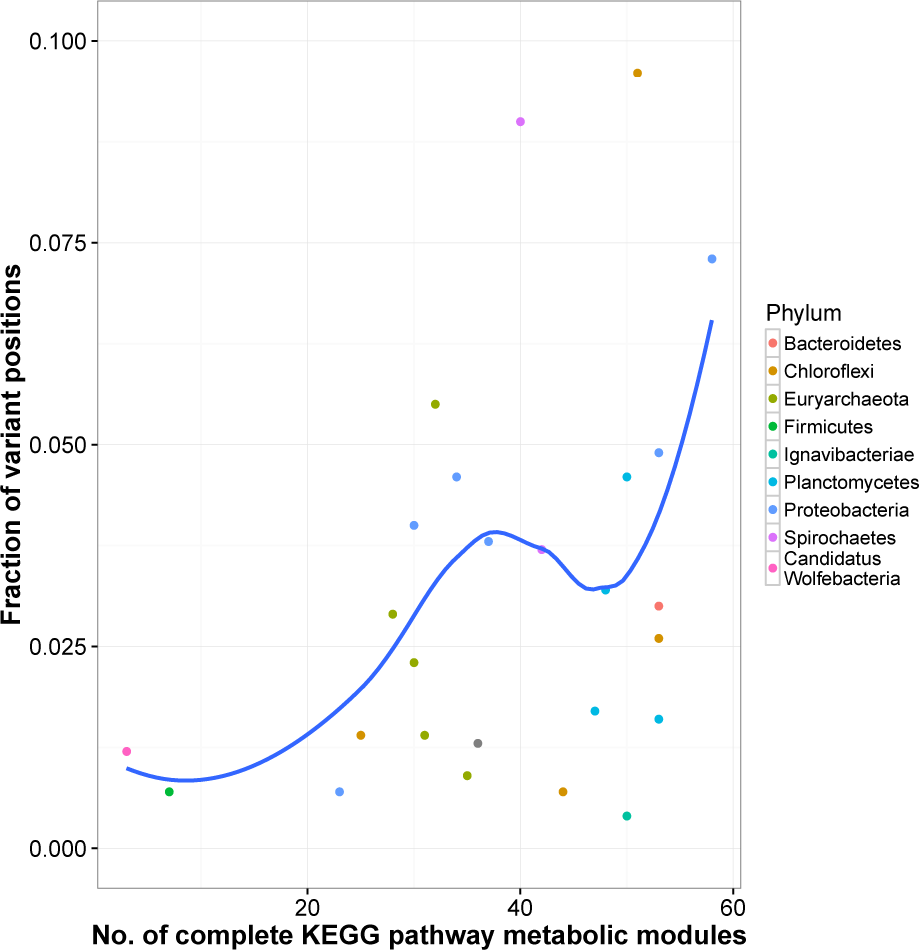
Variant frequency on SCGs against number of KEGG pathway modules for 26 AD MAGs. Variant frequency on the single-copy core genes for the 26 AD MAGs against the number KEGG metabolic pathways that were complete or near complete, defined as greater than 75% of a path through the module present. The curve shows a loess model fit to the data. A linear regression of variant frequency against module number gave (*R*^2^ = 0.1132,*p − value* = 0.05).

**Figure S14.**
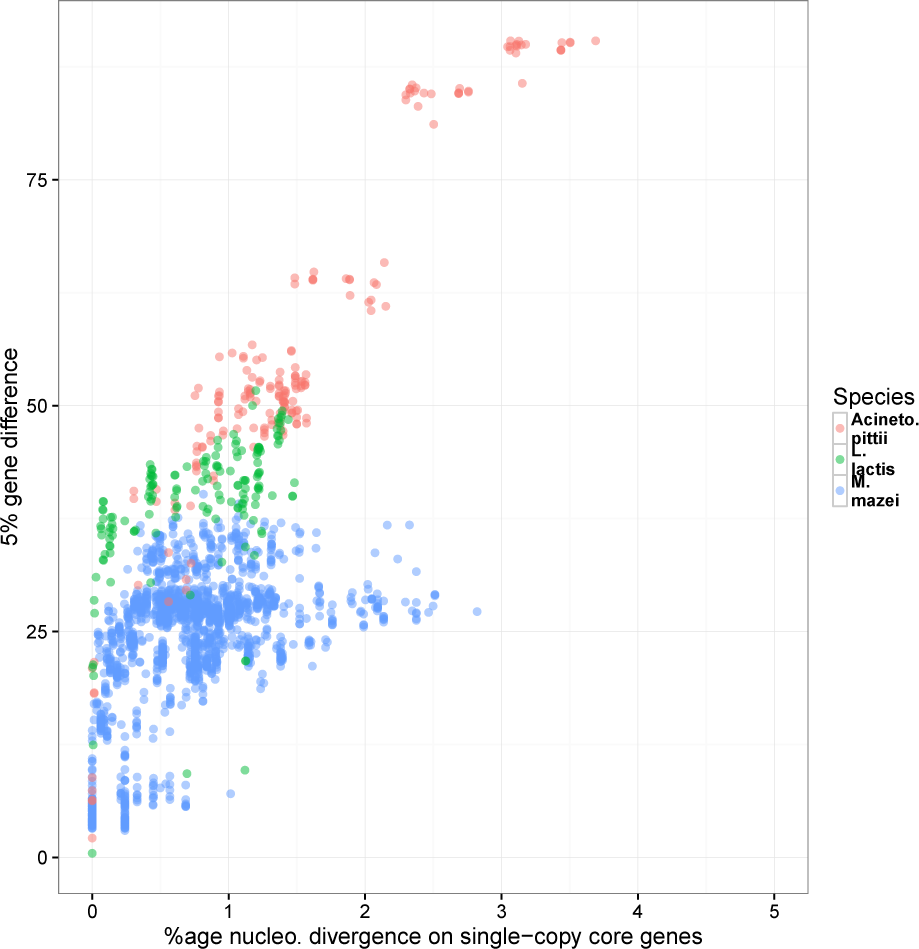
Comparison of nucleotide divergence on core genes with 5% gene cluster divergence for strains deriving from three environmental species. For each species (*Methanosarcina mazei*,*Lactococcus lactis* and *Acinetobacter pittii*), strains were downloaded from the NCBI list of bacterial and archaeal genomes. We then compared percentage nucleotide divergence on the core genes with percentage of 5% gene clusters shared between each pair of strains with nucleotide divergence *<* 5% for each species. In all three cases the relationship was highly significant but with different regression coefficients (*Methanosarcina mazei* — coeff. 7.7, Adjusted R-squared: 0.2246,*p − value <* 2.2*e −* 16,*Lactococcus lactis* — coeff. 8.2, Adjusted R-squared: 0.2401,*p−value* = 3.829*e*−11 and *Acinetobacter pittii* — coeff. 21.4, Adjusted R-squared: 0.8892, *p − value <* 2.2*e −* 16).

**Figure S15.**
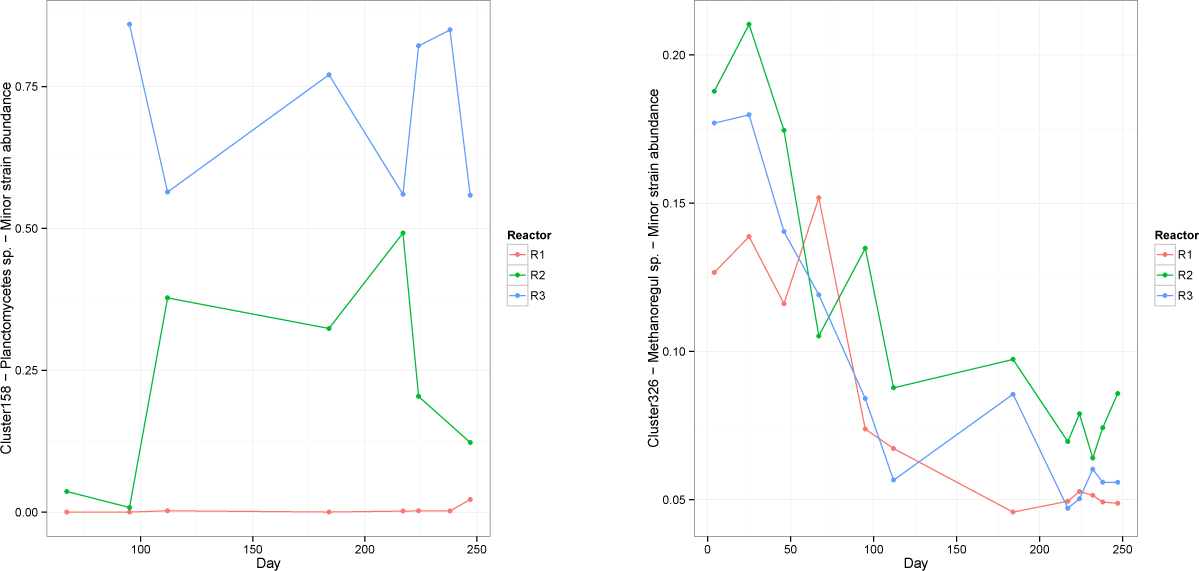
Strain abundance against time for two AD MAGs. Left) Cluster 158 - Planctomycetes sp. - abundance is shown in the bottom half of the three reactors against time Right) Cluster 326 Methanoregula sp.

## References

[1] Sanaa A. Ahmed, Joy Awosika, Carson Baldwin,Kimberly A. Bishop-Lilly, Biswajit Biswas, Stacey Broomall, et al. Genomic comparison of escherichia coli o104:h4 isolates from 2009 and 2011 reveals plasmid, and prophage heterogeneity, including shiga toxin encoding phage stx2. 7(11): 1–22, 11 2012.

[2] Mads Albertsen, Philip Hugenholtz, Adam Skarshewski, Kare L. Nielsen, Gene W.Tyson, and Per H. Nielsen. Genome sequences of rare, un-cultured bacteria obtained by differential coverage binning of multiple metagenomes. Nature Biotech., 31(6): 533+, JUN 2013.

[3] J Alneberg, BS Bjarnason, I de Bruijn, M Schirmer, J Quick,UZ Ijaz, L Lahti, N J Loman, A F Anderson, and C Quince. Binning metagenomic contigs by coverage and composition. Nat. Methods, 11:1144–1146, 2014.

[4] Yoav Benjamini andYosef Hochberg. Controlling the False Discovery Rate: A Practical and Powerful Approach to Multiple Testing. J. Roy. Statist. Soc. Ser. B, 57(1): 289–300, 1995.

[5] C M Bishop. Pattern Recognition and Machine Learning. Springer, 2006.

[6] S. Boisvert, F. Raymond, E. Godzaridis, F. Laviolette, and J. Corbeil. Ray Meta: scalable de novo metagenome assembly and profiling. Genome Biol., 13:R122, 2012.

[7] Christopher T. Brown, Laura A. Hug, Brian C. Thomas, Itai Sharon, Cindy J. Castelle, Andrea Singh, Michael J. Wilkins, Kelly C. Wrighton, Kenneth H. Williams, and Jillian F. Banfield. Unusual biology across a group comprising more than 15% of domain Bacteria. Nature, 523:208–211, 2015.

[8] James H Campbell, Patrick O’Donoghue, Alisha G Campbell, Patrick Schwientek, Alexander Sczyrba, Tanja Woyke, Dieter Söll, and Mircea Podar. UGA is an additional glycine codon in uncultured SR1 bacteria from the human microbiota. Proc. Natl. Acad. Sci. U.S.A., 110(14): 5540–5, apr 2013.

[9] A T Cemgil. Bayesian inference for nonnegative matrix factorisation models. Computational Intelligence and Neuroscience, page 785152, 2009.

[10] F.D. Ciccarelli, T. Doerks, C. von Mering, C.J. Creevey, B. Snel, and P. Bork. Toward automatic reconstruction of a highly resolved tree of life. Science, 311:1283–7, 2006.

[11] A. Corduneanu and C. M. Bishop. Variational Bayesian model selection for mixture distributions. In T. Jaakkola and T. Richardson, editors, Artificial Intelligence and Statistics 2001, pages 27–34. Morgan Kaufmann, 2001.

[12] Christopher J Creevey, Tobias Doerks, David A Fitzpatrick, Jeroen Raes, and Peer Bork. Universally distributed single-copy genes indicate a constant rate of horizontal transfer. PloS One, 6(8):e22099, jan 2011.

[13] A M Eren, Ö C Esen, C Quince, J H Vineis, H G Morrison, M L Sogin, and T O Delmont. Anvi’o: an advanced analysis and visualization platform for’ omics data. PeerJ, 3:e1319, 2015.

[14] F. Favero, T. Joshi, A. M. Marquard, N. J. Birkbak, M. Krzystanek, Q. Li, Z. Szallasi, and A. C. Eklund. Sequenza: allele-specific copy number and mutation profiles from tumor sequencing data. Ann Oncol., 26(1): 64–70, 2015.

[15] A Gelman, J B Carlin, H S Stern, D B Dunson, A Vehtari, and D B Rubin.Bayesian Data Analysis, Third edition. Chapman & Hall, 2013.

[16] Curtis Huttenhower, Dirk Gevers, Rob Knight, Sahar Abubucker, Jonathan H. Badger, Asif T.Chinwalla, et al. Structure, function and diversity of the healthy human microbiome. Nature, 486(7402): 207–214, JUN 14 2012.

[17] D. Hyatt, P.F. Locascio, Hauser L.J., and Uberbacher E.C. Gene and translation initiation site prediction in metagenomic sequences. Bioinformatics, 28:2223–2230, 2012.

[18] R. S. Kaas, C. Friis, D. W. Ussery, and F. M. Aarestrup. Estimating variation within the genes and inferring the phylogeny of 186 sequenced diverse escherichia coli genomes.BMC Genomics, 13:577, 2012.

[19] M. Katoh and M. Kuma. Mafft: a novel method for rapid multiple sequence alignment based on fast fourier transform. Nucleic Acids Res., 30:3059–3066, 2002.

[20] D.R. Kelley and S.L. Salzberg. Clustering metagenomic sequences with interpolated markov models. BMC Bioinform., 11:544, 2010.

[21] D D Lee and H S Seung. Algorithms for non-negative matrix factorization. Adv. Neural Inf. Process Syst., 13:556–562, 2001.

[22] Christophe Leys, Christophe Ley, Olivier Klein, Philippe Bernard, and Laurent Licata. Detecting outliers: Do not use standard deviation around the mean, use absolute deviation around the median. Journal of Experimental Social Psychology, 49(4): 764–766, 2013.

[23] Heng Li. A statistical framework for snp calling, mutation discovery, association mapping and population genetical parameter estimation from sequencing data. Bioinformatics, 27(21): 2987–2993, 2011.

[24] Heng Li andRichard Durbin. Fast and accurate long-read alignment with Burrows-Wheeler transform. Bioinformatics, 26(5): 589–595, 2010.

[25] N. J. Loman, C. Constantinidou, M. Christner, H. Rohde, J.Z. Chan, J. Quick, J.C. Weir, C. Quince, G.P Smith, J.R. Betley, M. Aepfelbacher, and M.J. Pallen. A culture-independentsequence-based metagenomics approach to the investigation of an outbreak of shiga-toxigenic escherichia coli O104:H4.J.A.M.A., 309:1502–10, 2013.

[26] N. J. Loman, J. Quick, and J. T. Simpson. A complete bacterial genome assembled de novo using only nanopore sequencing data. Nat. Methods, 12:733–735, 2015.

[27] C. Luo, R. Knight, H. Siljander, M. Knip, R. J. Xavier, and D. Gevers. Constrains identifies microbial strains in metagenomic datasets. Nature Biotech., page doi:10.1038/nbt.3319, 2015.

[28] Emilie E. L. Muller,Nicolás Pinel, Cédric C. Laczny, Michael R. Hoop-mann, Shaman Narayanasamy, Laura A. Lebrun, et al. Community-integrated omics links dominance of a microbial generalist to fine-tuned resource usage. Nat Commun, 5, 2014.

[29] Radford M. Neal. Markov chain sampling methods for Dirichlet process mixture models. J. Comp. Graph., 9:249–265, 2000.

[30] J. D. O’Brien, X. Didelot, Z. Iqbal, L. Amenga-Etego, B. Ahiska, and D. Falush. A bayesian approach to inferring the phylogenetic structure of communities from metagenomic data. Genetics, 3:925–37, 2014.

[31] Yu Peng, Henry C.M. Leung, S.M. Yiu, and Francis Y.L. Chin. Idbaud: A de novo assembler for single-cell and metagenomic sequencing data with highly uneven depth. Bioinformatics, 2012.

[32] PA Pevzner, H Tang, and MS Waterman. An eulerian path approach to DNA fragment assembly. Proc. Natl. Acad. Sci. U.S.A., 98:9748–9753, 2001.

[33] M.N. Price, P.S. Dehal, and A.P. Arkin. Fasttree 2 – approximately maximum-likelihood trees for large alignments. PLoS ONE, 5:e9490, 2010.

[34] Melanie Schirmer, Umer Z Ijaz, Rosalinda D’Amore, Neil Hall, William T Sloan, and Christopher Quince. Insight into biases and sequencing errors for amplicon sequencing with the illumina miseq platform. Nucleic acids res., gku1341, 2015.

[35] M Scholz, D V Ward, E Pasolli, T Tolio, M Zolfo, F Asnicar, D T Truong, A Tett, A L Morrow, and N Segata. Strain-level microbial epidemiology and population genomics from shotgun metagenomics. Nature Met., page doi:10.1038/nmeth.3802, 2016.

[36] N Segata, L Waldron, A Ballarini, V Narasimhan, O Jousson, and C Huttenhower. Metagenomic microbial community profiling using unique clade-specific marker genes. Nat Methods, 9:811–4, 2012.

[37] I. Sharon, M.J. Morowitz, B.C. Thomas, E.K Costello, D.A. Relman, and J.F. Banfield. Time series community genomics analysis reveals rapid shifts in bacterial species, strains, and phage during infant gut colonization. Genome Res., 23:111–20, 2013.

[38] Y. Wang, H.C. Leung, S.M. Yiu, and F.Y. Chin. Metacluster 5.0: a two-round binning approach for metagenomic data for low-abundance species in a noisy sample. Bioinformatics, 28:i356–i362, 2012.

[39] M Welling andM Weber. Positive tensor factorization. Pattern Recognition Letters, 22:1255–1261, 2001.

[40] O. Zagordi, A. Bhattacharya, N. Eriksson, and N. Beerenwinkel. Shorah: estimating the genetic diversity of a mixed sample from next-generation sequencing data. BMC Bioinform., 12:119, 2011.

